# High Enzyme Promiscuity in Lignin Degradation Mechanisms in *Rhodopseudomonas palustris* CGA009

**DOI:** 10.1101/2025.02.15.638337

**Authors:** Mark Kathol, Niaz Bahar Chowdhury, Cheryl Immethun, Adil Alsiyabi, Dianna Morris, Michael J. Naldrett, Rajib Saha

## Abstract

Lignin is a universal waste product of the agricultural industry and is currently seen as a potential feedstock for more sustainable manufacturing. While it is the second most abundant biopolymer in the world, most of it is currently burned as it is a very recalcitrant material. Many recent studies, however, have demonstrated the viability of biocatalysis to improve the value of this feedstock and convert it into more useful chemicals, such as polyhydroxybutyrate, and clean fuels like hydrogen and n-butanol. *Rhodopseudomonas palustris* is a gram-negative bacterium which demonstrates a plethora of desirable metabolic capabilities, including aromatic catabolism useful for lignin degradation. This study uses a multi-omics approach, including the first usage of CRISPRi in *R. palustris*, to investigate the lignin consumption mechanisms of *R. palustris*, the essentiality of redox homeostasis to lignin consumption, elucidate a potential lignin catabolic superpathway, and enable more economically viable sustainable lignin valorization processes.

## Introduction

Lignin is famously known as the second most abundant biopolymer on earth, consisting of about 20-30% of all plant biomass^1–3^. Lignin is a universal waste product from every biorefinery industry which works with lignocellulosic biomass feedstocks, with Kraft pulping paper manufacturing as the most common source. Around 118,000,000 metric tons were produced globally in 2015 from this process alone^4^. Instead of being utilized in other downstream biorefineries as a feedstock, it is commonly burned^5^ as production costs for valorizing lignin are prohibitively high, or cannot compete with cheaper, non-sustainable methods of production^6,7^. Many recent studies have demonstrated that lignin can be converted into other more useful products such as itaconic acid^8^, carbon nanotubes^9^, and various aromatic intermediates like benzene^10^ and arenes^11,12^.

Therefore, a very large potential market exists for utilizing this waste product should it be made economically viable. Lignin, when depolymerized, is converted to a range of aromatic phenylpropanoids, with slightly differing chemical structures, the main difference being the amount of meta-methoxylation present on the aromatic ring. H lignin has no methoxy groups, G lignin possesses strictly one, and S lignin has both meta positions methoxylated. These chemicals are known as lignin breakdown products (LBPs). Because of this variation, a large portion of studies which focus on lignin valorization often investigate specific microorganisms for their use of promiscuous enzymes, which are able to react with a range of LBPs indiscriminately. Such microorganisms include *Comamonas testosteroni*^13–15^, *Pseudomonas putida* KT2440^16,17^, *Sphingobium* sp. SYK-6^18,19^, and of particular interest in this study, *Rhodopseudomonas palustris* CGA009 (hereafter *R. palustris*)^20,21^. *R. palustris* is an incredibly metabolically versatile organism that is known for being able to catabolize various aromatic substrates, such as LBPs^22–25^. *R. palustris* is a common soil bacterium which is capable of all four modes of metabolism, namely photoheterotrophy, photoautotrophy, chemoheterotrophy, and chemoautotrophy, and possesses both carbon dioxide and nitrogen fixation abilities^26^. It can also produce valuable commodities such as vanillic acid^27^, polyhydroxybutyrate, a bioplastic precursor^21^, and fuels like hydrogen^28^ and n-butanol^29^,

The main strategy to increase the economic viability of bacteria as lignin valorization chassis is to utilize metabolic engineering and tailor strains to maximize titer and yield of bioproducts. However, in order to employ this method, the metabolism of lignin breakdown products must be characterized. *R. palustris*’ metabolism of certain H lignin units has been verified previously^25^, but its aerobic and anaerobic metabolism of G and S units are not as well understood. *R. palustris* possesses a wide variety of redundant and promiscuous enzymes, which can make deciphering catabolic pathways challenging. In this study, we utilize a multi-omics approach to understand the mechanism by which each LBP is consumed (Fig. 1). Next-gen-sequencing, transcriptomics, and label-free quantitative proteomics are utilized in tandem in this study to determine the enzymes associated with each reaction and screen for potential metabolic pathways. The quantification of various metabolites and CRISPRi are then used to verify our proposed pathway. This study will enable future lignin valorization products which utilize this versatile organism to increase their economic viability.

**Fig 1.**
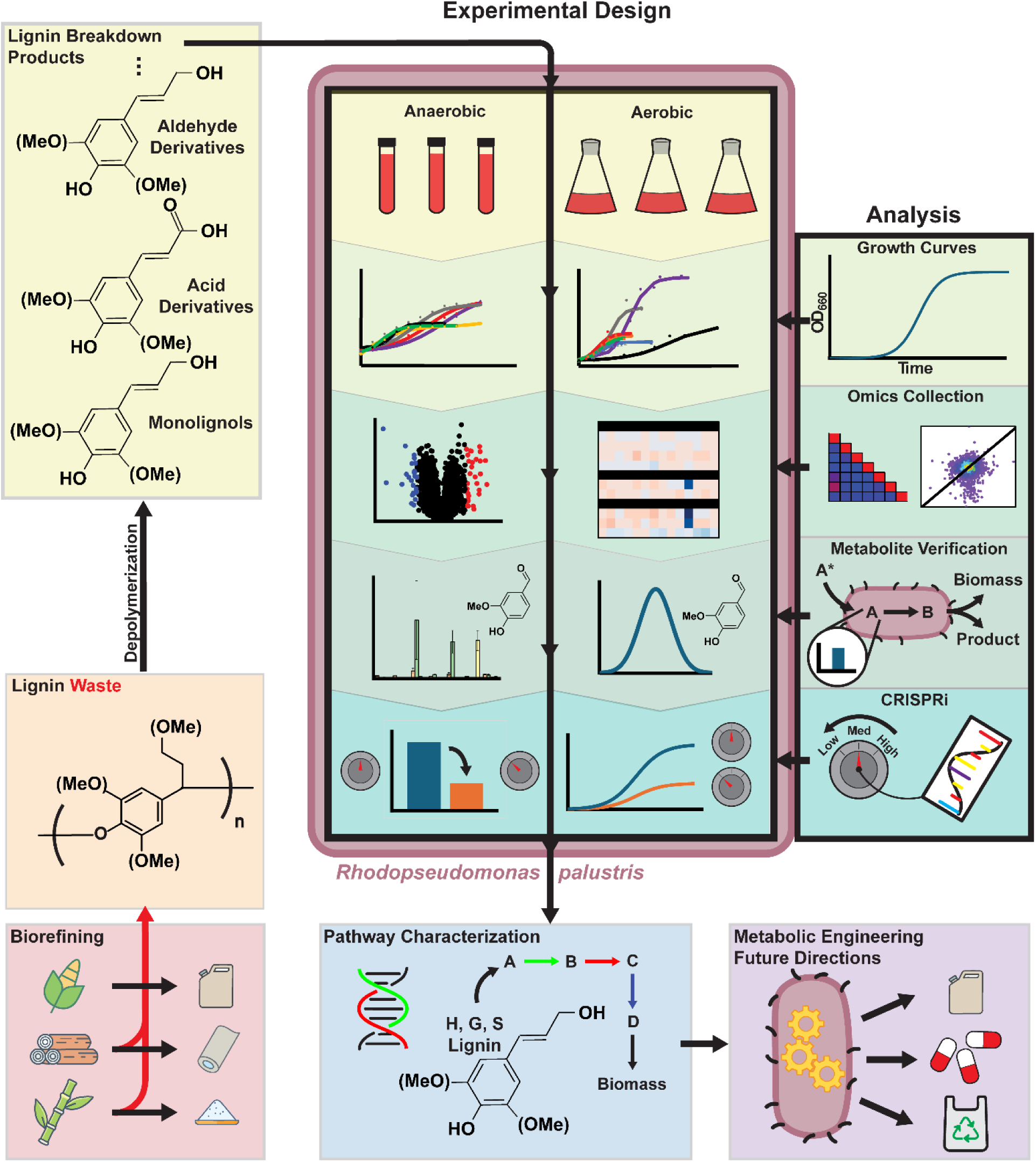
Workflow.

## Results and Discussion

### *R. palustris* Growth on LBPs

To establish an initial understanding of *R. palustris*’ ligninolytic abilities, wild-type *R. palustris* cultures were provided with a variety of LBPs as well as kraft lignin as their main carbon source. Due to lignin’s amorphous structure between plants, the abundance of individual LBPs can vary drastically from differing feedstock materials^3^. To narrow down the list of possible LBPs which *R. palustris* may consume, the monolignols *p-*coumaryl alcohol, coniferyl alcohol, and sinapyl alcohol, as well as two acid derivatives, *p-*coumarate and sodium ferulate, were investigated as carbon substrates (Fig. 2a&b). These substrates were chosen due to their abundance in kraft lignin depolymerization^3,30^. The purpose of this initial experiment was to determine under which conditions *R. palustris* can catabolize this selection of carbon substrates, as well as growth rates and maximum cell concentrations. *R. palustris* was also grown with acetate to serve as a control due to its simple catabolism. We then generated growth curves for each carbon source under both aerobic and anaerobic conditions (Fig. 2c&d) (Supplementary Data 1). From our growth experiment, *R. palustris* can notably utilize each type of lignin as a carbon source.

**Fig. 2:**
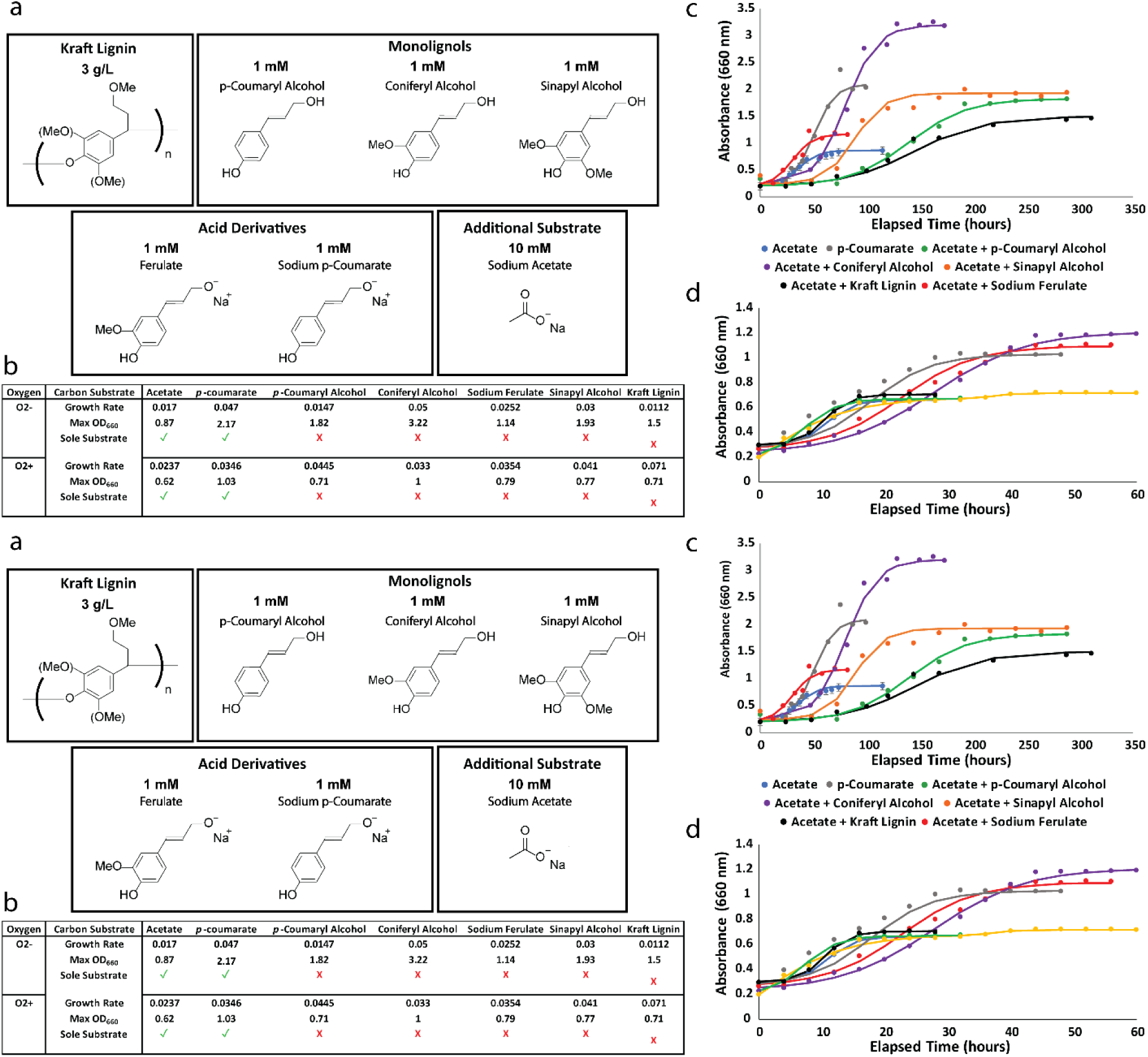
Wild type *R. palustris* growth on various LBPs. **(a)** Carbon substrates used in this study, consisting of commonly abundant lignin depolymerization products including monolignols and acid derivatives. **(b)** Growth statistics for WT *R. palustris* when supplemented with 1mM of corresponding carbon substrate. **(c)** Anaerobic and **(d)** aerobic growth curves of WT. *R. palustris* on selected carbon substrates

*R. palustris* was conditionally co-fed acetate when it was unable to consume the LBP as a sole carbon substrate, though generally when supplemented with an LBP, *R. palustris* grows to a higher concentration than our acetate controls. *p-*coumarate appears to be readily consumed by *R. palustris* without acetate supplementation under both aerobic and anaerobic conditions, while *p-*coumaryl alcohol and the meta-methoxylated carbon sources, coniferyl and sinapyl alcohol, are more recalcitrant. Sinapyl alcohol in particular depicts the most interesting growth behavior, as there appears to be two distinct exponential growth curves, which is characteristic of diauxic growth. Diauxic growth occurs when a bacterium selectively consumes one of two carbon substrates in solution at a time, consuming the more readily metabolized substrate first, which given the simplicity of acetate metabolism, is almost certainly acetate. Another stark trend from our growth curves is that *R. palustris* grows to much higher concentrations under anaerobic growth. This suggests that either photophosphorylation is a more efficient method of generating ATP than oxidative phosphorylation, or that some enzymes along the pathway may be oxygen labile. *R. palustris* appears to at least partially consume each LBP under at least one of the oxic conditions. Anaerobic growth with coniferyl alcohol fed as a carbon substrate produces the highest average OD_660_ of 3.19 ± 0.005, while three other carbon substrates, *p*-coumarate, *p*-coumaryl alcohol, and sinapyl alcohol, appear to converge at roughly the same max OD_660_ of 1.80 - 2.10.

### Transcriptomics and Proteomics Data Analysis

Given that our initial experiments showed definitive evidence of *R. palustris*’ ability to at least partially catabolize these LBPs into biomass, we investigated the genes and associated proteins which are involved with various aromatic catabolic pathways which *R. palustris* possesses. Genes and proteins that are upregulated in samples containing LBPs compared to the control acetate samples can therefore be correlated with monolignol and broader aromatic consumption. Using this methodology, *R. palustris*’ catabolism route of *p-*coumarate, one of the selected carbon substrates for this study, has already been discovered^25^. To assess the activity of each of the aromatic pathways held by *R. palustris* when supplemented with each LBP individually, we first gathered gene lists from previous annotation of *R. palustris* and catabolic pathways for the degradation of multiple aromatic compounds. This list includes genes for the degradation of *p-* coumarate, protocatechuate, homogentisate, benzoate, phenylacetate, and a meta-cleavage pathway^26^. After a list of enzymes for the complete catabolic pathways was generated, we obtained transcriptomic and proteomic datasets for *R. palustris* cultures grown under our LBP conditions (Supplementary Data 2 & 3). This was to determine which pathway would be most upregulated, and therefore likely responsible for LBP catabolism under each condition, as well as indicate any enzyme promiscuity between sample groups containing differing LBPs (Fig. 3). From our datasets, we filtered expression and abundance values, performed quantile normalization, and obtained Log_2_ fold changes and p*-*values for gene transcripts and proteins comparative to our acetate controls for all samples (Fig. 3 a&b). To obtain an initial understanding of the similarity in protein regulation for differing LBP catabolisms, we constructed correlation triangular heatmap matrices, which contain Pearson correlation values between the sample groups (Fig. 3 c&d). A very interesting trend emerges from these heatmaps, as a moderate correlation exists on average between the aerobic sample groups (∼0.27), with some stark outliers having very high correlation, e.g. *p-*coumarate/coniferyl alcohol (0.62) and *p-*coumarate/sinapyl alcohol (0.79). However, a much higher correlation on average is present for the anaerobic samples, (∼0.46), with the same outliers (0.82 and 0.70 respectively). This strongly suggests that the same metabolic pathways may be employed by *R. palustris* for the consumption of all lignin types regardless of meta-methoyxlation.

**Fig 3:**
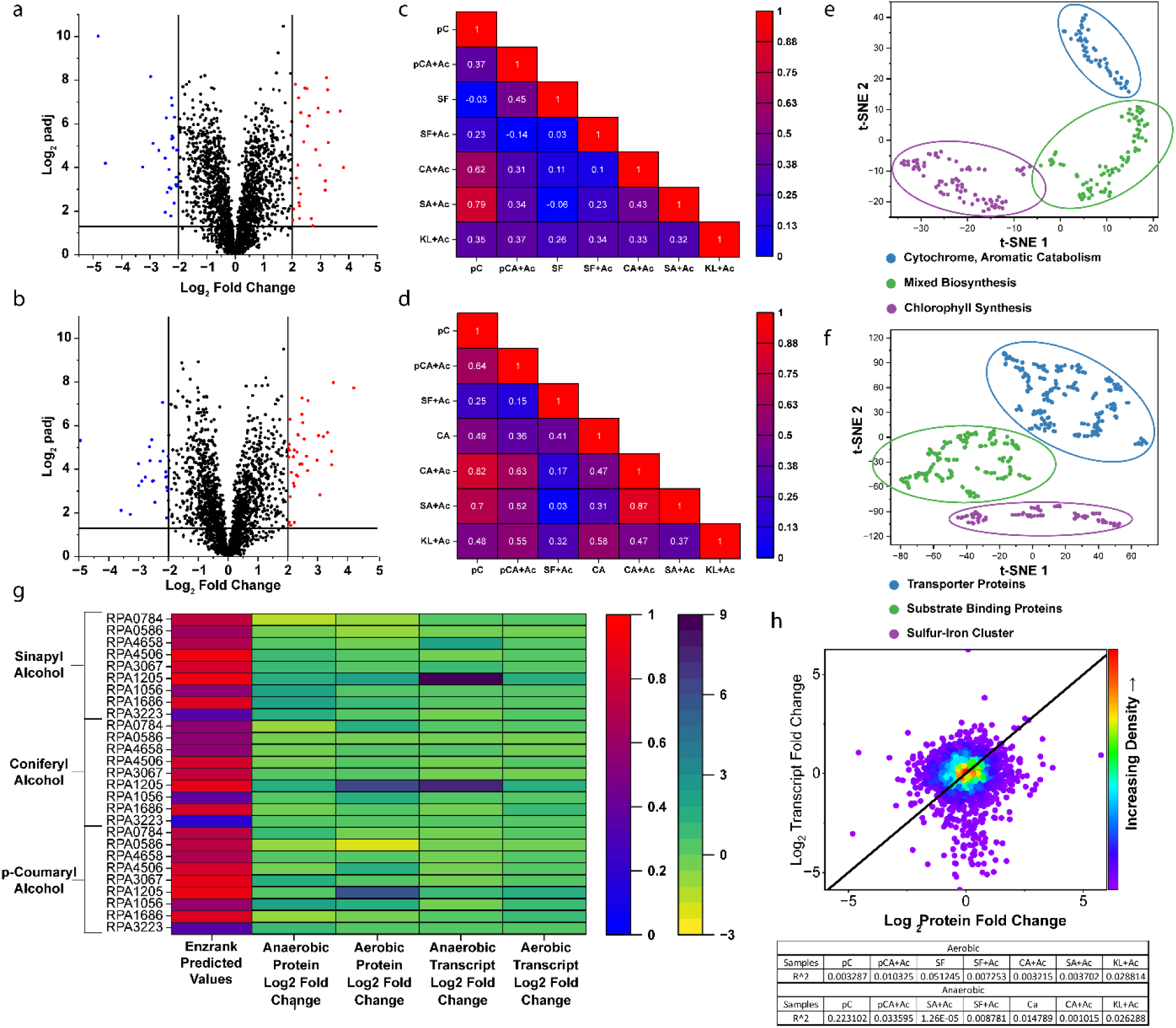
Summary of *R. palustris* combined omics analysis. **(a)** Volcano plot of upregulated aerobic *p-*coumarate vs acetate proteome. **(b)** Volcano plot of upregulated aerobic *p-*coumarate vs acetate proteome. **(c)** Correlation between upregulated proteome profiles between aerobic and **(d)** anaerobic LBP growth conditions. **(e)** t-SNE of top 200 upregulated features across all aerobic conditions and (f) top 400 proteins across all anaerobic growth conditions. **(g)** Combined Heatmap depicting EnzRank predicted enzyme-substrate binding values for various upregulated alcohol dehydrogenases and their correlative transcriptomic and proteomic Fold2 upregulation values. **(h)** Correlation between transcriptome and proteome for *p-*coumarate, with table depicting R^2 values for other aerobic and anaerobic cultures.

A commonly used tool when analyzing large datasets to determine if patterns exist between different measured variables is principal component analysis (PCA). T-distributed stochastic neighbor embedding (t-SNE) is a variation of PCA which essentially performs the same function but is able to account for some of the non-linear relationships which may exist in gene expression and protein abundance. To determine any other significant protein activity outside of LBP catabolism, we also performed a t-SNE of the top 200 and 400 upregulated proteins in our aerobic and anaerobic samples respectively (Fig. 3e&f). Aerobically, the most dominant protein groups in each cluster were for cytochrome proteins, various biosynthesis pathways including polyhydroxybutyrate, and most interestingly, bacteriochlorophyll synthesis. The presence of cytochrome and bacteriochlorophyll proteins imply that one of the most important metabolic functions under any ligninolytic growth condition appears to be redox homeostasis. As LBPs are far more oxidized carbon sources compared to other substrates such as glucose and even acetate, a large portion of the proteome in each of these conditions provides reducing power to consume these substrates. This reconfirms findings from several studies that suggest that ligninolysis is not rate-limited thermodynamically but is instead constrained by the need to maintain redox homeostasis ^21,31–33^. Bacteriochlorophyll is very unexpected given that these cultures were grown aerobically but may imply that *R. palustris* may prefer to utilize photophosphorylation as a more efficient method of generating ATP when consuming oxidized compounds. Anaerobically, the most prominent clusters consist of transporter proteins, various substrate-binding proteins, and iron-sulfur clusters. Considering that under most anaerobic conditions *R. palustris* grows to much higher ODs (Fig. 2c), this would imply that photosynthesis is a more efficient electron sink in terms of biomass yield per unit of substrate. The expected iron-sulfur protein cluster is present in these results which is consistent with anaerobic photoheterotrophic growth, and more of the proteome is dedicated to aromatic catabolic reactions than under the aerobic conditions, as seen with the transporter and substrate-binding proteins. The presence of the bacteriochlorophyll proteins under most of our aerobic conditions also indicates that *R. palustris* may have photosystem I active under all conditions.

In many organisms, certain proteins perform the same enzymatic function to ensure redundancy and enhance environmental fitness^34–36^. However, this can lead to ambiguities when trying to determine which enzymes are most active under specific conditions. For example, *R. palustris* has multiple non-specific alcohol and aldehyde dehydrogenases that are upregulated in several of our samples. This creates a problem when reconstructing metabolic pathways as, due to the redundancy, the enzyme responsible for performing a possible substrate-specific reaction can be confused with other enzyme of the same reaction class. To resolve these ambiguities, we used EnzRank^37^, a machine learning-based model that predicts the likelihood of enzymes binding to specific substrates. This helped narrow down the list of enzymes likely responsible for carrying out these reactions (Fig. 3 g). *R. palustris* may also employ multi-level regulation between transcriptome and proteome, as from our datasets, as our data show that only a few transcriptomes and proteomes exhibit even moderate correlation (Fig. 3h).

Lastly, to corroborate the information we have obtained from our omics analysis, we have also generated heatmaps detailing the state of upregulation for the compiled aromatic catabolic pathways. This was done to elucidate which are the most upregulated under each growth condition supplemented with our selected aromatics (Fig. 4). We have found that under almost every growth condition, the *p-*coumarate pathway appears to be the most upregulated proportional to its size. This, in light of the strong correlation that exists between both aerobic and anaerobic samples (Fig. 3h), leads us to believe that the *p-*coumarate enzymes act promiscuously among the carbon substrates, catalyzing the same reaction amongst the metabolite analogs independent of oxic conditions. The amount of promiscuity from this investigation hints at a possible superpathway, where similar carbon substrates are being funneled towards aromatic cleavage and the TCA cycle using the same set of enzymes^38^. Given that each of the lignin units have similar phenolic structure with varying amounts of meta-methoxy functional groups, we hypothesize that the enzymes responsible for *p-*coumarate metabolism have highly promiscuous activity with G and S lignin units. Using this information, we have created a superpathway diagram that outlines all the proposed enzymes, reaction types, and metabolites for each necessary reaction (Fig. 5). These pathways, whether operating aerobically or anaerobically, direct all LBPs toward the TCA cycle through a series of extensive reduction reactions.

**Fig 4:**
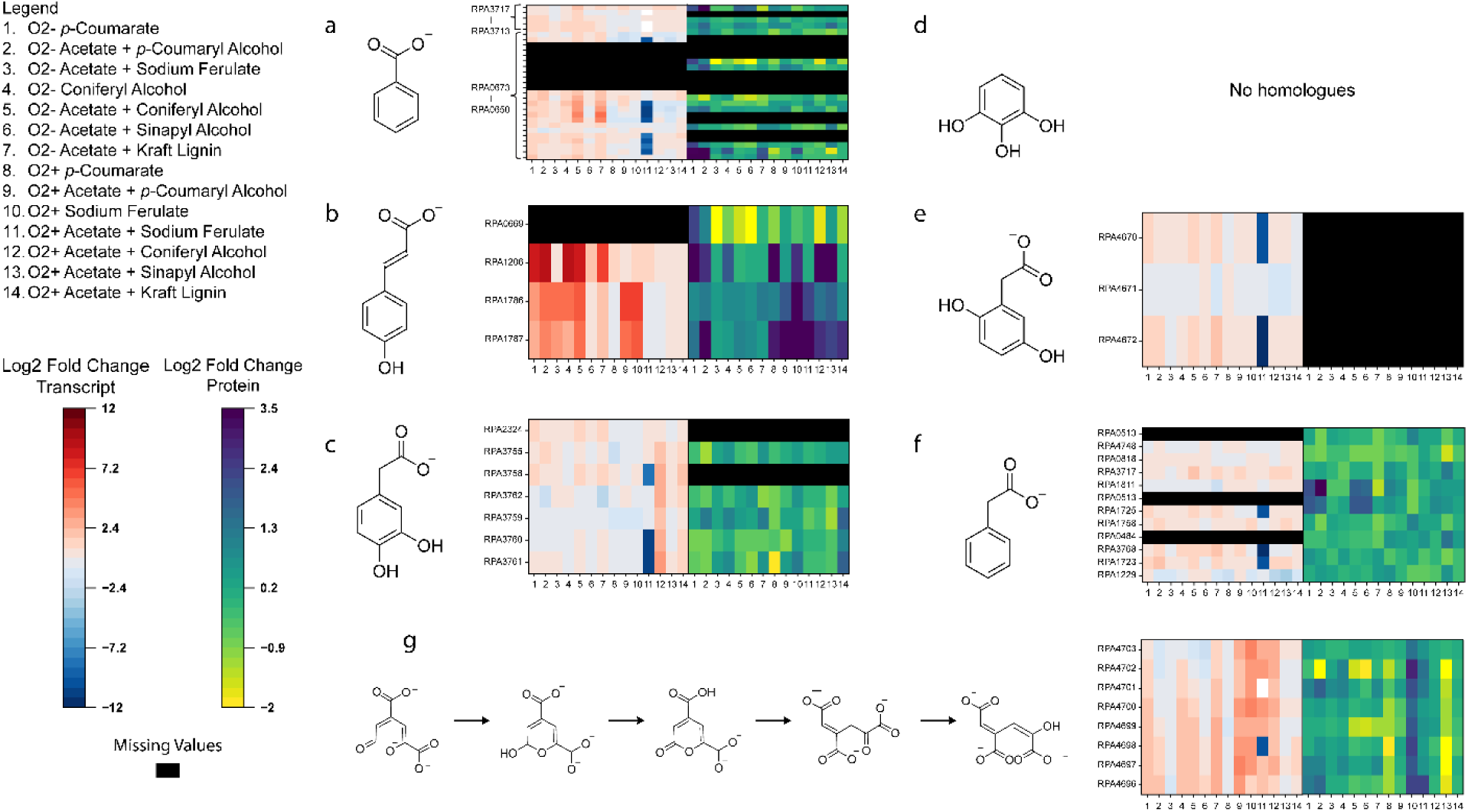
Upregulation states of common, homologous, aromatic catabolic pathways. Upregulation states for **(a)** benzoate, **(b)** *p-*coumarate, **(c)** protocatechuate, **(d)** pyrogallol, **(e)** homogentisate, **(f)** phenylacetate, and **(g)** meta-cleavage homologous catabolic pathways in *R. palustris*. Missing values are marked in black as not all transcripts or proteins abundancies are always captured during data collection.

**Fig 5:**
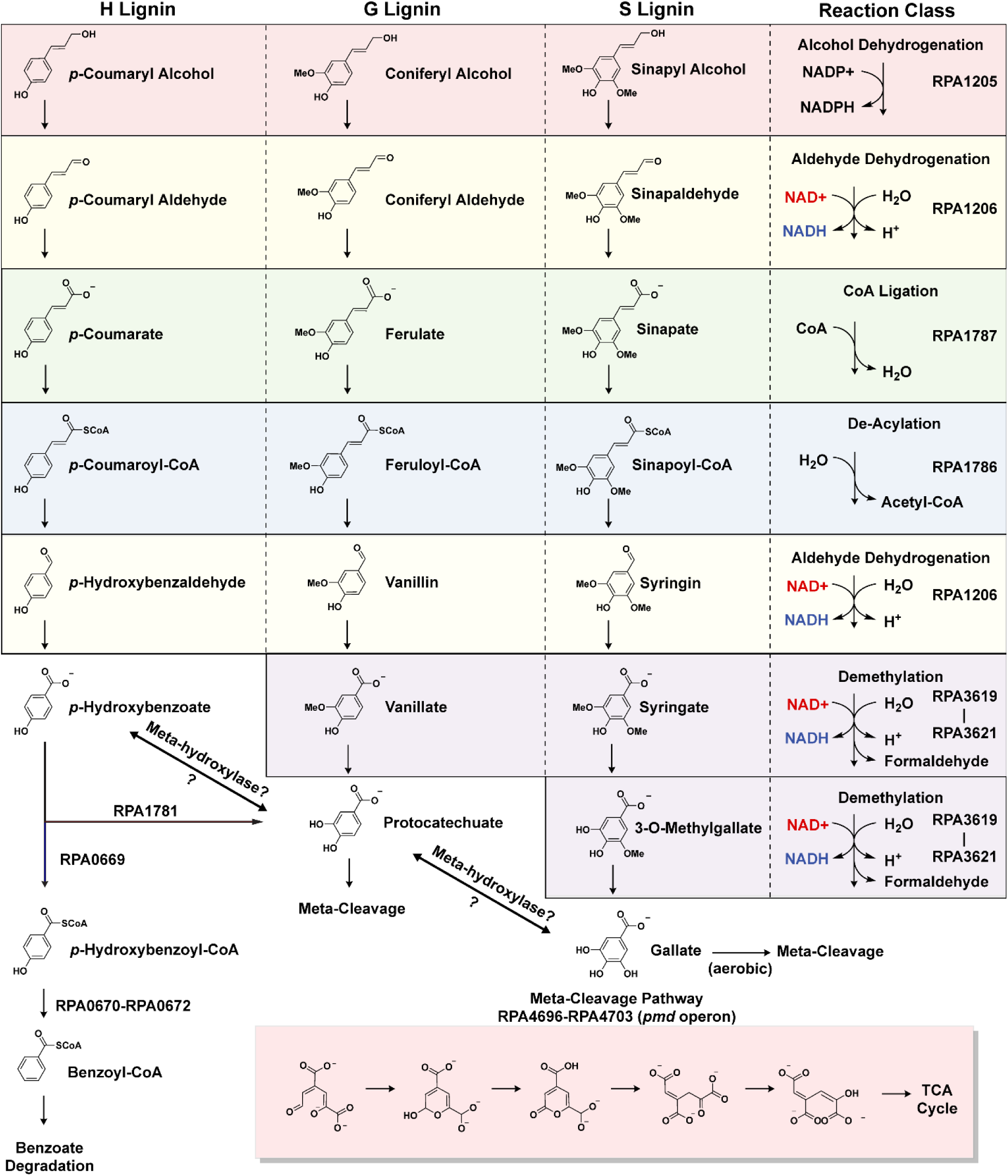
Proposed catabolic superpathway for lignin breakdown products by wild type *R. palustris* CGA009. Breakdown of monolignol aromatics using *p-*coumarate reactions and meta-cleavage.

### Alcohol and Aldehyde Dehydrogenation, Deacetylation, and Redundancy

At the beginning of the superpathway are the monolignol substrates, *p-*coumaryl, coniferyl, and sinapyl alcohol. These substrates are typically reduced to their acid derivatives using both alcohol and aldehyde dehydrogenase enzymes. From our transcriptome and proteome profiles, many enzymes for both classes were upregulated. ^34–37^ Of the upregulated enzymes, EnzRank determined *rpa1205*, an alcohol dehydrogenase which is simultaneously the most upregulated enzyme of its class across all conditions, is most likely to catalyze the monolignols to aldehyde derivatives. EnzRank also determined *rpa1206*, an aldehyde dehydrogenase responsible for the catalysis of *p-*hydroxybenzaldehyde to 4-hydroxybenzoate in the *p-*coumarate catabolic pathway, most likely converts the monolignol aldehyde derivatives to acid conjugates, *p-*coumarate, ferulate, and sinapate (Fig 3. g). A de-acetylation reaction then converts these compounds by removing the acetyl-CoA group from the alkyl chain via *rpa1786*, reducing these substrates to *p-* hydroxybenzaldehyde, vanillin, and syringin. These, next, undergo subsequent reduction to their respective acid derivatives by utilizing *rpa1206* again, which also corroborates results found by Zhang et al.^27^, where several aldehyde dehydrogenases were deleted from *R. palustris*’ genome, rpa1206, rpa1687, and rpa1725, to optimize vanillin production. An aldehyde dehydrogenase activity heatmap can be found in Supplementary Fig. 1.

### Vanillate and Syringate Demethylation

After being reduced from alcohol to acid derivatives and undergoing subsequent de-acylation, H, G, and S lignin each require distinct metabolic steps to undergo aromatic cleavage. The resulting H lignin metabolite *p-*hydroxybenzoate, under anaerobic conditions, is directly converted into *p-* hydroxybenzoyl-CoA and subsequently to benzoyl-CoA. The G and S lignin unit metabolite analogues, however, protocatechuate, and sinapate respectively, possess one and two meta-methoxy functional groups (Fig. 5). Currently, there is no known aromatic ring cleavage pathway which includes these methoxy functional groups, and other organisms such as *Acinetobacter baylyii* ADP1^39^, *Pseudomonas* sp. HR199^40^, and *Pseudomonas putida* KT2440 commonly demethylate these methoxy groups by conversion to protocatechuate from vanillate, and 3-O-methylgallate or gallate from sinapate^41^. *R. palustris* contains homologous protein subunits to *Pseudomonas putida*’s *VanAB* as *rpa3619* and *rpa3621*, which are currently annotated as putative vanillate O-demethylase subunits A and B respectively. A study performed by Oshlag et al. noted that the *rpa3619-rpa3621* was essential for S lignin unit catabolism, interestingly even under anaerobic conditions, despite the reaction class of this enzyme typically involving dioxygen^23^. In this study anaerobic *R. palustris* cultures were observed upregulating both *rpa3619-rpa3621* transcripts and proteins aerobically and anaerobically. This study, as a result, observed that *R. palustris* was capable of the demethylation of vanillate and gallate even under anaerobic conditions, concluding that these reactions must not require oxygen. Given that these transcripts are exclusively, significantly upregulated (*P* < 0.05, Log_2_ Fold Change > 2) both aerobically and anaerobically for G and S lignin units, we propose that this enzyme set catalyzes the conversion of vanillate and sinapate to protocatechuate, and 3-O-methylgallate and gallate respectively. *VanAB* is also known to be oxygen labile^23^, which may explain the higher ODs under anaerobic growth when G and S lignin is supplemented to *R. palustris*, as the vanillate demethylase compromised enzyme activity could cause bottlenecking issues under aerobic growth.

### Aerobic Aromatic Ring Cleavage

At this point in the superpathway, the aromatic rings in protocatechuate and gallate must be cleaved to be further processed into biomass (Fig. 5). Aerobically, protocatechuate can be produced from the catabolism of both H and G lignin, as *R. palustris* contains an enzyme homologous to *Pseudomonas putida*’s *p-*hydroxybenzoate hydroxylase (*rpa1781*). This enzyme aerobically converts p-hydroxybenzoate into protocatechuate, which can then undergo aerobic cleavage. Aromatic rings are enzymatically cleaved through three main pathways, the 2,3 meta cleavage, the ortho cleavage, and the 4,5 meta cleavage pathways^13^. The primary method by which bacteria degrade aromatic compounds is through ortho-cleavage of the ring, also known as the beta-ketoadipate pathway^42^. However, this pathway is not present in *R. palustris*. *R. palustris* does, however, possess 4,5 meta-cleavage in the form of a *pmd* operon. The upregulation of the *pmd* operon shows mixed results in both our aerobic and anaerobic samples.

In the aerobic state, transcripts for the *pmd* operon are predominantly upregulated in samples containing p-coumaryl alcohol, sodium ferulate, and coniferyl alcohol. In contrast, transcript levels are upregulated for p-coumarate, p-coumaryl alcohol, coniferyl alcohol, and kraft lignin samples in the anaerobic state. However, protein abundance data reveal a very different pattern, with little to no significant upregulation of the *pmd* operon proteins (*P* > 0.05, Log2 Fold Change < 1). Furthermore, the upregulation observed in the *pmd* operon transcripts is less pronounced compared to the strong upregulation seen in the *p*-coumarate pathway genes. This does not necessarily imply that the meta-cleavage pathway is not involved in aromatic ring cleavage in *R. palustris*. Despite the lack of upregulation, it is possible that this section of ligninolysis does not present a bottleneck. For now, we hypothesize that the *pmd* operon’s 4,5 meta-cleavage pathway is utilized for the catabolism of protocatechuate and gallate, thereby contributing to the resolution of aerobic lignin catabolism.

### Anaerobic Aromatic Ring Cleavage

So far there exists very few known pathways for anaerobic degradation and cleavage of aromatic compounds, the most prominent of which is the benzoyl-CoA degradation pathway. *R. palustris* can catabolize H lignin units anaerobically through the conversion of *p-*coumaric acid to 4-hydroxybenzoate, to further be converted to benzoyl-CoA which is then consumed through the BAD pathway^25^. However, compared to *p-*hydroxybenzoate degradation, anaerobic G and S lignin catabolic pathways in any organism have not been as thoroughly annotated. Proposed pathways include removing the meta-hydroxyl groups to catalyze gallate to protocatechuate, and protocatechuate to *p-*hydroxybenzoate^25^, as well as several unfinished annotated pathways involving trans-hydroxylation and CoA-ligation of G lignin by *Thauera aromatica*^43^. The removal of hydroxyl groups from these compounds would funnel the metabolites towards the benzoate degradation pathway, yet in our own omics analysis and in a previous study^23^, the BAD pathway is not upregulated in any anaerobic samples containing S lignin. Benzoate degradation is interestingly upregulated in the case of anaerobic coniferyl alcohol and *p-*coumarate consumption, but not for sodium ferulate or *p-*coumaryl alcohol samples. This leaves seemingly paradoxical evidence that *R. palustris* consumes selectively H lignin acid derivatives and G lignin alcohols through the BAD pathway, but not H lignin alcohols or G lignin acid derivatives, even though there should be no lack of enzymatic capabilities in consuming these compounds similarly^23^. Another possible method particularly for S lignin consumption is the decarboxylation of gallate to pyrogallol, such as that which occurs in various gut bacterium such as *Eubacterium oxidoreducens* G41^44^, soil bacterium such as *Rhodococcus opacus*^45^, and phylogenetically similar soil bacteria such as *Bradyrhizobium japonicum*^46^, and *Pelobacter acidigallici*^45^. This is also unlikely as *R. palustris* does not contain enzyme homologs to documented pyrogallol catabolic pathways. There also could be a strict dehydroxylation of S lignin units after demethylation. Meta-dehydroxylation has also not been studied well in any organism, as the enzyme class for this reaction does not exist. Most confusingly is that the *pmd* operon is upregulated in certain anaerobic samples, even when compared to their aerobic analogues (Fig. 4g) (Supplementary Fig. 2). This pathway is expected to be inactive under anaerobic conditions because it requires dioxygen. However, similar sample preparations were used to maintain anaerobic conditions, as in a previous study ^23^. Culture tubes were filled completely leaving no headspace, and N_2_ was used to purge air from prepared samples, yet the *pmd* operon is upregulated for our anaerobic sodium ferulate samples and partially for the coniferyl alcohol samples. More research will be needed to verify how the *pmd* operon can function under anaerobic conditions, but a similar phenomenon may be occurring for anaerobic ring cleavage as with anaerobic vanillate and syringate demethylation. One last possibility that remains is that *R. palustris* possesses a novel anaerobic method of ring cleavage specific to both G and S lignin, or contains unique catabolic pathways with respect to both, such as that found within *Thauera aromatica42,* which to the best of our knowledge is completely unannotated. Considering that these pathways are currently not annotated, it is not possible to perform a comparative genomic analysis and subsequent comparative upregulation analysis of these pathways. For now, however, we postulate that a combination of the meta-cleavage and BAD pathways are utilized to cleave H, G, and S lignin units anaerobically.

### Pathway Confirmation

To provide additional supporting evidence to our proposed superpathway and determine if the individual metabolites are present in solution, we performed a targeted metabolite analysis of our *R. palustris* samples (Supplementary Data 4). We chose to measure a select group of metabolites to determine if *R. palustris* is capable of performing specific enzymatic steps along our proposed pathway. Amongst our selected metabolites we measured *p-*hydroxybenzoate, vanillate, and syringate to determine if *R. palustris* can perform the *p-*coumarate pathway steps up to and including the de-acylation step. *p-*hydroxybenzoate, vanillate, and syringate are also archetypal degradation metabolites respective to the H, G, and S lignin types respectively. We also measured protocatechuate and gallate for the purposes of determining if *R. palustris* is capable of performing the demethylation step along the pathway. Benzoate and protocatechuate were also measured as surrogates for the activity of both the BAD pathway and meta-cleavage due to the difficulty of obtaining standards for intermediate metabolites. *R. palustris* cultures were harvested at mid-exponential when incubated under both aerobic and anaerobic conditions supplied with each of the LBPs consistent with our previous omics and growth experiments (Fig. 6 a&b). Based on our proposed pathway, we should see specific metabolites under each growth condition (Fig. 6c). Not all of the predicted metabolites may be present under each growth condition, as some reactions present in the pathway may have much higher rates than others, which would prevent accumulation in those cases.

**Fig 6:**
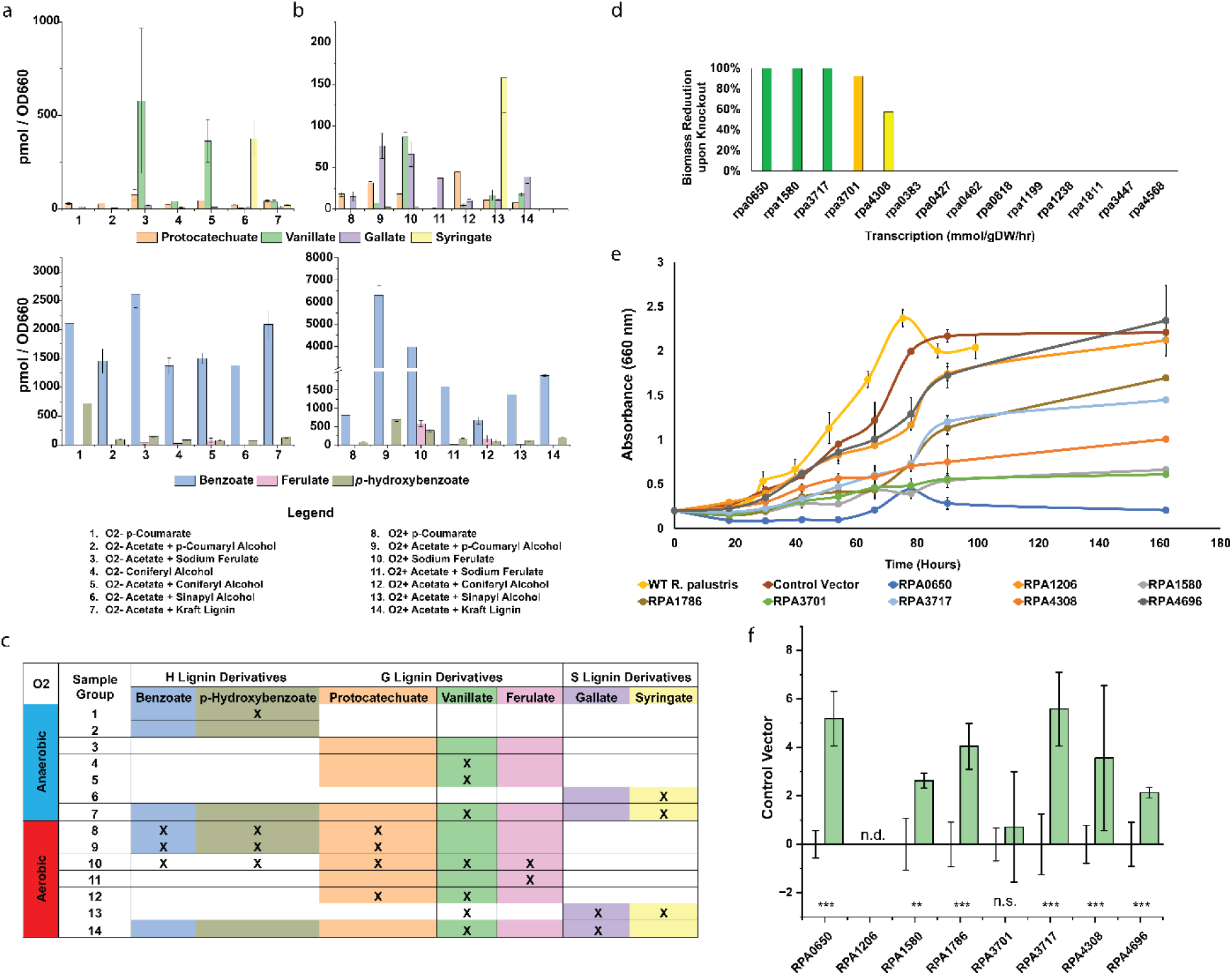
Metabolite confirmation and redox homeostasis CRISPRi knockdown. **(a**) Anaerobic and **(b)** aerobic metabolite profiles of LBP fed wild-type *R. palustris* cultures. **(c)** Expected metabolites under each growth condition. Highlighted cells represent predicted presence based on the proposed catabolic superpathway. X represents actual significant accumulation in the corresponding sample. **(d)** Growth reduction predictions based on essentiality of various NAD+ synthases. **(e)** Anaerobic photoheterotrophic growth curves of CRISPRi repressed strains supplemented solely with 1mM *p-*coumarate. **(f)** RNA-based CRISPRi repression rates for corresponding CRISPRi constructs (n = 4, ** = P < 0.005, *** = *P* < 0.0005, nd = not detected, ns = no significance). Error bars without corresponding value bars represent one standard deviation of control vector expression rates.

The presence of metabolites under each condition is largely consistent with our proposed pathway (*P* < 0.05), with some noteworthy trends. *p*-Hydroxybenzoate appears in nearly all of our samples including our wild-type acetate controls, both aerobically and anaerobically, at a baseline concentration of around 100 pmol/OD. This is likely due to the supplementation of our bacterial culturing medium, PM, with *p*-aminobenzoate, which can be easily converted to p-hydroxybenzoate. *p*-aminobenzoate is commonly supplemented into growth media as it is a precursor to folic acid, which is vital for many metabolic processes. However, significant concentrations of *p*-hydroxybenzoate are observed in the aerobic *p*-coumaryl alcohol (3046 pmol/OD) and sodium ferulate (542 pmol/OD) samples, while anaerobically, it is only significantly present in the *p*-coumarate samples (722 pmol/OD). The fact that *p*-hydroxybenzoate and benzoate are present during aerobic growth could be indicative of aerobic consumption of H lignin through the BAD pathway, though its absence in various anaerobic samples contrarily indicate that the BAD pathway is not utilized for these other carbon substrates. Vanillate profiles reveal a more interesting trend. Significant vanillate concentrations are observed as expected in G lignin samples, including aerobic coniferyl alcohol (5 pmol/OD) and sodium ferulate (85 pmol/OD), but accumulate to much higher levels under the anaerobic conditions, such as coniferyl alcohol (364 pmol/OD) and sodium ferulate (579 pmol/OD). Intriguingly, vanillate also appears in our aerobic sinapyl alcohol cultures (16 pmol/OD). Syringate, as anticipated, is exclusively found in S lignin samples, both aerobic (158 pmol/OD) and anaerobic (376 pmol/OD), however, the lack of accumulation of gallate in even our S lignin cultures suggests that either the subsequent cleavage step occurs much faster which would prevent accumulation, or that syringate is instead converted to protocatechuate. Overall, the metabolite profile up to the de-acylation step is consistent with the proposed pathway, thus supporting its accuracy.

As with p-hydroxybenzoate, benzoate concentrations were also elevated at a high baseline (1500 pmol/OD) in all of our samples, likely due to the addition of *p*-aminobenzoate to the culture media. Benzoate remained significantly present in our aerobic *p*-coumaryl alcohol (6306 pmol/OD) and sodium ferulate (3976 pmol/OD) samples. However, the anaerobic benzoate profiles did not follow the same pattern. Interestingly, benzoate concentrations were unexpectedly higher (2726 pmol/OD) in our control acetate samples than in any other sample group, complicating trend analysis. Despite this, the aerobic benzoate profiles, in combination with the p-hydroxybenzoate concentrations, suggest that *R. palustris* may be channeling both *p*-coumaryl alcohol and sodium ferulate toward the benzoate pathway under aerobic conditions, even though previous studies ^24,47^ have shown that CGA009 cannot utilize benzoate aerobically. Given that benzoate is present in much higher concentrations (by a factor of 10) than other metabolites, it’s possible that *p*-hydroxybenzoate and benzoate are accumulating without being fully consumed, with the remaining carbon from *p*-coumaryl alcohol and sodium ferulate directed toward the meta-cleavage pathway as runoff. Additionally, the presence of *p*-hydroxybenzoate and benzoate in our aerobic sodium ferulate samples, along with vanillate in the aerobic sinapyl alcohol samples, provides strong evidence for meta-hydroxylase activity in *R. palustris*. However, our anaerobic *p*-coumaryl alcohol samples depict a distinct lack of any desired measurable metabolite. This, when paired with the fact that *R. palustris* grows slowly anaerobically with *p*-coumaryl alcohol, suggests that metabolites are prevented from accumulating to any significant amount.

For G lignin metabolites, protocatechuate was significantly present in nearly all of our aerobic samples (Fig. 6 a&c) which suggests that *R. palustris* funnels all LBPs through 4,5 meta-cleavage generally when grown aerobically. This also suggests meta-hydroxylase activity as protocatechuate concentrations are significant even in our aerobic S lignin sample group. Our anaerobic acetate culture controls were too unreliable for the measurement of protocatechuate, making protocatechuate trends difficult to interpret. So far, the metabolite profiles almost completely agree with the proposed pathway including the demethylation step.

With most of the metabolites for our proposed superpathway confirmed, to supplement verification of the proposed superpathway, the activity of each of the enzymes needs to be confirmed. This can be done traditionally with a complementation study, or other biochemical assays such as an *in vitro* protein activity assay, though our efforts to generate a knockout mutant for various genes in *R. palustris* has been met with mixed success.

### Essential Genes Associated with Redox Homeostasis

Since oxygen is not available as an electron sink under anaerobic conditions, there remains the possibility that the growth rate, thereby the catabolism of LBPs, by *R. palustris* may be limited by a lack of NAD+ regeneration and internal redox balance. Hence, to find alternate electron sinks, we used our recently published metabolic and expression (ME-) model of *R. palustris*^48^, where *p-* coumarate was used as the carbon source and ammonium was used as the nitrogen source. At first, all the reactions that produced NAD+ were sequentially turned off in the ME-model. Out of 29 reactions that generate NAD+, the ME-model did not produce any biomass growth rates for 20 reactions. We later collected all the involved genes for those 29 reactions and found that 14 genes were involved in the 29 reactions (Supplementary Data 5). We sequentially turned off each gene in the ME-model to see its impact on the biomass growth rate (Fig. 6d). We found, by turning off *rpa3717* (enoyl-CoA hydratase), *rpa1580* (proline dehydrogenase), and *rpa0650* (cyclohex-1-ene-1-carboxyl-CoA hydratase), the ME-model did not produce any biomass growth rate. Moreover, turning *rpa3701* (5,10-methylenetetrahydrofolate reductase) and *rpa4308* (phosphoglycerate dehydrogenase) off reduced the biomass growth rate by 92% and 57% respectively. Thereby, *rpa3717*, *rpa1580*, *rpa0650*, *rpa3701*, and *rpa4308* can be the alternate electron sources that reproduces NAD+. Further

To test the ME-model predictions, we performed CRISPRi experiments by knocking down each of the genes individually. A CRISPRi expression system was designed using a previously optimized plasmid for heterologous protein expression^49^, and sgRNA sequences were incorporated for RPA0650, RPA1580, RPA3701, RPA3717, and RPA4308. We also developed two more CRISPRi plasmids to confirm the essentiality of the aromatic catabolic enzymes *rpa1786,* a hydratase, and *rpa4696*, a *p-*oxalomesaconate tautomerase. These two enzymes were chosen for knockdown as they are non-redundant enzymes associated with two separate mechanisms in lignin degradation. *rpa1787* was selected to determine its essentiality in the initial reduction and de-acylation section of lignin metabolism, while *rpa4696* was chosen to determine if meta-cleavage was still occurring anaerobically and is vital for lignin consumption even without the presence of oxygen.

We incubated each of the CRISPRi strains of *R. palustris* anaerobically with 1 mM of p-coumarate (Fig. 6e). The resulting growth curves revealed some intriguing and significant trends, particularly regarding the impact of knocking down genes encoding NAD+ generating enzymes, which had a severely detrimental effect on growth. The maximum change in OD_660_ for the NAD+ encoding gene knockdown strains compared to wild-type *R. palustris* showed a progressive reduction: *rpa3717* (71.08%), *rpa4308* (49.55%), *rpa1580* (32.52%), and *rpa3701* (29.90%), with *rpa0650* exhibiting nearly no growth (10.13%). In contrast, knockdowns of the catabolic enzymes *rpa1786* and *rpa4696* showed little to no reduction in growth, suggesting these genes do not significantly impact overall growth. These results strongly indicate that lignin catabolism in *R. palustris* is not rate-limited by the abundance of catabolic enzymes, but rather by the availability of NAD+, which is essential for the metabolic processes involved.

To confirm the repression of each of the mRNA transcripts, we performed RT-qPCR with primers designed to bind to each of the repressed enzymes respective transcripts. Using this method, we were able to measure repression for each of the target mRNA transcripts with exception to RPA1206 (Fig. 6f). Several primer sets were used to determine the relative abundance of RPA1206 transcripts, however, each either failed to produce amplification or did not have an acceptable primer efficiency. Our CRISPRi expression system was able to reliably repress RPA0650 (Average Log_2_ Fold Repression of 5.18 ± 1.12), RPA1580 (2.63 ± 0.31), RPA 1786 (4.04 ± 0.94), RPA3717 (5.58 ± 1.52) RPA4308 (3.56 ± 2.99) and RPA4696 (2.13 ± 0.22). RPA3717 was unfortunately not significantly repressed using our dCas9 expression system. Given that, of the confirmed repressed transcripts the genes associated with redox-homeostasis produced the most growth inhibition, this further cements the idea that redox-homeostasis is more important than catabolic enzyme abundance in lignin catabolism.

## Conclusions

The lignin challenge remains to be resolved as the overwhelming majority of the waste lignin produced today is still burned without further application. *R. palustris*, for its known ability to consume a wide variety of aromatics, is a promising candidate to serve as a biotechnological chassis to convert lignin towards products that are otherwise produced non-sustainably. In this study, we propose that the enzymes *R. palustris* utilizes for the catabolism of *p-*coumarate may also be used for the catabolism of other lignin types. Our omics analysis consists of the comparison of upregulation states of *R. palustris* when fed with LBP substrates, and corroboration of predictive tools such as EnzRank to resolve enzyme ambiguities. Through our omics analysis, we also determine that a combination of the benzoate and meta-cleavage pathways may be utilized by *R. palustris* to consume each of the lignin substrate types. Even more notably, meta-cleavage may be active in some anaerobic conditions. As confirmation of our proposed pathway, metabolite profiles were then measured to verify *R. palustris* ability to perform certain enzymatic reactions. This study also utilized metabolic modelling to determine where the bottleneck for lignin catabolism occurs, which appears to be in maintaining redox homeostasis, rather than the abundance of enzyme or thermodynamic limitation. To verify this, our study, to our knowledge, demonstrated the first application of CRISPRi dCas9 repression in *R. palustris*. Utilizing our CRISPRi dCas9 expression system, we verified that the knockdown of various NAD+ genes severely reduced growth. Lastly, to complete the verification of each of the enzymes associated with our superpathway, a complementation study, or similar will need to be performed.

Further studies involving 13C-labelled carbon substrates and multiple metabolite sampling points during exponential growth can grant further insight into where bottlenecks can be occurring along the pathway, as well as experimentally determined metabolic fluxes. These were not performed as the labelled substrates were not commercially available. Once these bottlenecks are known, modifications to the proteins mentioned here can be made to further increase ligninolytic throughput for *R. palustris*. The CRISPRi vector developed in this study could also be used to further probe gene function and essentiality under other desirable growth conditions. Another hypothesis worth investigating is the possibility that fatty acid and polyhydroxybutyrate production “compete” for reducing power during ligninolysis. Through the use of dCas9 or Cas9 to repress or outright remove a gene from *R. palustris*, we can determine if the removal of either polyhydroxybutyrate or lipid biosynthesis increases flux directed towards the other. Our findings, in summary, can guide future ligninolysis studies with *R. palustris* to transform it into a metabolic engineering chassis, more effectively valorize lignin, and create more sustainable production methods for a variety of chemicals.

## Materials and Methods

### Strain Growth Conditions

*Rhodopseudomonas palustris* (Molisch) van Niel BAA-98 CGA009 was obtained from the American Type Culture Collection (ATCC). All strains used in this study are described in Supplementary Table 1 and were stored at −80°C. NEB® 10-beta Competent *Escherichia coli* was used for all plasmid construction, followed by transformation into Stellar™ Competent dam^-^/dcm^-^ *Escherichia coli* (ATCC) to remove demethylation before insertion into *R. palustris*. All *R. palustris* strains were stored in a final concentration of 20% (v/v) glycerol while all E. coli strains were stored with 15% (v/v) glycerol. *R. palustris* and E. coli strains retrieved from storage were first grown on solid 112 Van Niel’s (VN) media and solid LB media (Miller, AMRESCO) respectively with appropriate antibiotic^49^.

Seed cultures of *R. palustris* were grown aerobically before conducting LBP growth experiments in 50 mL of Photosynthetic Media (PM) in 250 mL Erlenmeyer flasks supplemented with 20mM NaC_2_H_3_O_2_, 10 mM HCO_3_ and 15.2 mM (NH_4_)_2_SO_4_^50^. Seed cultures were diluted to OD_660_ = 0.2 in 50 mL of PM in 250 mL Erlenmeyer flasks if grown aerobically, or 13.5 mL of PM in 14 mL round bottom tubes with no headspace, at 30°C and 275 rpm. Resazurin dye was also used as an oxygen indicator in order to confirm anaerobic conditions. All aerobic and anaerobic cultures were supplemented with 10 mM HCO_3_ and 15.2 mM (NH_4_)_2_SO_4,_ as well as the appropriate LBP at a final concentration of 1mM and 10mM. All growth curves of our *R. palustris* cultures are included in Supplementary Data 1.

### Strain Construction

All plasmids were constructed with a protocol as described previously^49,51^. All oligonucleotides were purchased through Eurofins Genomics or Integrative DNA Technologies™. The sequences of all genes used in this study are provided in Supplementary Table 2. All plasmids in this study used synthetic biology parts, of which the performance and effectiveness when employed in *R. palustris* has been defined previously according to Immethun et al. 2022, with exception to CRISPR dCas9^51^. Briefly, PCR was conducted using Phusion Hot Start II Polymerase (Thermo Scientific™). Upon verifying the correct size of the amplicon through gel electrophoresis with 1x TAE buffer 1% agarose gels, PCR product was then purified with the Monarch® DNA Gel Extraction Kit (New England Biolabs® Inc). E. coli was incubated overnight at 30°C and 275 rpm in 4 mL of LB media in 14 mL round bottom tubes. E. coli cultures were then diluted to 1/40^th^ OD_600_ in 4mL of LB media and incubated for 1 hour at 30°C and 275 rpm until mid-exponential growth phase, and washed at room temperature according to previous literature^52^. After washing, cells were then transformed with assembled hot fusion or blunt end ligation product through electroporation at 2.5 kV. Electroporated E. coli cells were then incubated in 0.450 mL of LB media at 30°C and 275 rpm for 1.5 hours before streaking onto LB plates supplemented with 30 μg/mL kanamycin, and subsequent incubation at 30°C overnight. Colonies were then selected from the plates and grown in 4 mL of fresh 30 μg/mL kanamycin supplemented LB media overnight. All E. coli cultures were then stored at −80°C at a final concentration of 15%(v/v) glycerol. Constructed plasmids from E. coli cultures were harvested using the Purelink™ Quick Plasmid Miniprep Kit (Invitrogen™).

Whole plasmid sequencing was performed on each extracted plasmid to ensure that all plasmids were correctly assembled after transformation. After extraction, plasmids were then transformed into dam^-^/dcm^-^ *Escherichia Coli* in order to obtain demethylated plasmid to improve retention rate. Demethylated plasmids were then extracted and used to transform wild-type *R. palustris*. *R. palustris* was transformed using the same protocol as described, with the following exceptions.

*R. palustris* was incubated in VN liquid and solid media instead of LB for all incubation steps. *R. palustris* was grown in VN media for 18 hours prior to dilution to OD_660_ = 0.2 in fresh VN media, before additional incubation overnight to OD_660_ = 0.6 or mid-exponential for transformation. After electroporation and streaking onto VN plates supplemented with 300 μg/mL kanamycin, *R. palustris* was then incubated at 37°C for up to 1 week after transformation as colonies take far longer to appear due to *R. palustris*’ longer doubling time. Colonies from *R. palustris* were then streaked onto new kanamycin supplemented VN plates and incubated for approximately 5 days before growth in 300 μg/mL VN liquid media. *R. palustris* cultures were then stored at −80°C in a final concentration of 20% (v/v) glycerol. For more detailed step by step instruction on strain growth conditions and construction, please see Kathol et al., 2023^53^.

### RNA Extraction and RT-qPCR

sgRNA sequences for dCas9 plasmid constructs were designed using the online tool CHOPCHOP (https://chopchop.cbu.uib.no/). *R. palustris* strains containing CRISPRi expression vectors were diluted to OD_660_ = 0.2 at 30°C and 275 rpm, and induced with 1mM IPTG. RNA from dCas9 containing *R. palustris* cultures was extracted using a modified TRIZOL method^53^. RNA degradation quality was determined via bleach gel as described in Aranda et al.^54^. RNA extracts were then reverse-transcribed using the Multiscribe® Reverse Transcriptase protocol. RT-qPCR primers were designed and tested as previously described. RT-qPCR primer set information and efficiencies, as well as cDNA concentrations for each amplicon are outlined in Supplementary Table 4. Primer concentrations for RT-qPCR were initially evaluated using Go Taq^®^ Master mix, with primer concentrations ranging from 350-50 nM. A second set of PCR reactions without and DNA template were also prepared. These PCR reactions were then performed with the following settings, 95°C for 2 min, 40 cycles of (95°C for 45 s, 60°C for 45 s, 72 for X seconds—based on amplicon length), and then 72°C for 5 min. Each reaction was then run on 2% agarose gels to ensure a lack of undesirable PCR products.

Primer efficiencies for each primer set were determined using the same protocol as described previously^55^. Briefly, qPCR reactions were prepared with a 5x serial dilution of wild type *R. palustris* cDNA (500 – 0.0061 ng/μL). These reactions were then run with an Eppendorf Mastercycler Realplex with the following settings: 50°C for 2 minutes, 95°C for 2 minutes, followed by 40 cycles of 95°C for 15 seconds and 60°C for 1 minute, then followed by a melting curve, 95 °C for 15 seconds, 60°C for 15 seconds, then ramping to 95°C at a rate of 1.75°C/minute. Ct values obtained from these reactions were then plotted against the Log_10_ transformation of the cDNA concentration. The slope of this line then determined the primer efficiency, and primer efficiencies between 90-110% were accepted for determining relative transcript abundance. qPCR was then performed on four biological replicates with two technical duplicates to ensure consistent Ct reporting. Log_2_ fold change expression values were then calculated via ΔΔC_t_ method, with 16SrRNA serving as the housekeeping gene.

### Transcriptomic, Proteomic, Metabolite Data Acquisition

After constructing *R. palustris* LBP growth curves, mid-exponential phases were determined to harvest cultures for submission to various core facilities to obtain transcriptomic, proteomic, and metabolomic profiles. 2 replicates of each aerobic and anaerobic growth condition were grown and were treated with RNA*later* before submission to CD genomics. 5 replicates of each growth condition were collected and submitted to the UNL Metabolomics and Proteomics Core Facility for obtaining proteomic profiles, as well as an additional set of 3 replicates for collecting metabolite concentrations.

To obtain transcriptomic profiles, firstly, ribosomal RNA depletion was performed using NEBNext® rRNA Depletion Kit (Bacteria, New England Biolabs cat#E7860S). The rRNA-depleted RNA was purified by 2x RNAClean XP beads (Beckman Coulter) and eluted in 45ul of nuclease-free water. The purified RNA was then mixed with 4ul of NEBNext First Strand Synthesis Reaction Buffer and 1ul of random primers. The reaction was incubated at 94C for 12 mins for fragmentation. Subsequently, to perform first strand cDNA synthesis, 8ul of NEBNext Strand Specificity Reagent and 2ul of NEBNext First Strand Synthesis Enzyme Mix was added to the reaction and the reaction was incubated at 25C 10 mins, 42C 30 mins, 70C 15 mins. Second strand reaction was then performed by adding 8ul of NEBNext Second Strand Synthesis Reaction Buffer with dUTP Mix (10X), 4ul of NEBNext Second Strand Synthesis Enzyme Mix, and 48ul of nuclease-free water. The reaction was incubated at 16C 1hr. The cDNA was purified by 1.8x SPRIselect Beads (Beckman Coulter) and eluted in 50ul of nuclease-free water. The reaction was incubated at 16C 1hr. The cDNA was purified by 1.8x SPRIselect Beads (Beckman Coulter) and eluted in 50ul of nuclease-free water. Subsequently, endprep reaction was performed by adding 7ul of NEBNext Ultra II End Prep Reaction Buffer and 3ul of NEBNext Ultra II End Prep Enzyme Mix into 50 ul purified cDNA. Endprep reaction was incubated at 20C 30 mins and 65C 20 mins. Adaptor ligation reaction was then performed by adding 1ul of NEBNext Ligation Enhancer, 30ul of NEBNext Ultra II Ligation Master Mix, and 2.5ul of NEBNext Adaptor, diluted to 0.5uM in Adaptor Dilution Buffer. The mix was incubated at 20C 15 mins. 3ul of USER Enzymer (New England Biolabs) was then added to the ligation product and the reaction was incubated at 37C 15 mins. The ligated product was purified by SPRIselect Beads (Beckman Coulter) and eluted in 15ul of nuclease-free water. PCR was carried out by adding 25ul of NEBNext Ultra II Q5 Master Mix, 5ul of i5 Primer, and 5ul of i7 Primer into 15ul of purified ligated product. PCR was performed at 98C 30 seconds, 15 cycles of 98C 10 seconds and 65C 75 seconds, and a final extension at 65C 5 mins. The final library was then purified by SPRIselect Beads and loaded into Illumina NovaSeq6000 paired end 150 bp mode for sequencing.

### Label-free proteomics analysis

The cell pellets were lysed in Pierce RIPA buffer (Thermo Fischer Scientific, Waltham, MA, USA) containing 5 mM DTT and 1x protease inhibitor (complete EDTA-free protease inhibitor cocktail; Roche, Indianapolis, IN, USA) by shaking them at 95°C for 10 min on a thermomixer. The samples were then centrifuged at 16,000 x g for 15 min, and the supernatants were transferred to a new tube. The proteins were assayed using the CB-X protein assay (G-Biosciences, St Louis, MO, USA). 50 µg of reduced protein was alkylated with 20 mM iodoacetamide for 40 min and quenched with DTT. The proteins were then precipitated with acetone and the pellets washed 3 times with 70% ethanol. Proteins were resuspended in 50 µL of 50mM Tris/HCl, pH 8.0 containing 1 µg Lys-C and digested for 4 h, followed by further digestion with 1 µg trypsin overnight at 37°C. A quality control reference sample was prepared by mixing all the samples with the same 1:1 ratio to run between every 16 samples to check for instrument performance deviation. The sequence order of the samples was randomized using block randomization.

Each digest was run by nano liquid chromatography-tandem mass spectrometry (nanoLC-MS/MS) as previously described using an Ultimate 3000 RSLCnano system coupled to an Orbitrap Eclipse mass spectrometer (Thermo Fisher Scientific). Briefly, peptides were first trapped and washed on a trap column (Acclaim PepMap™ 100, 75µm x 2 cm, Thermo Fisher Scientific). Separation was then performed on a C18 nano column (Acquity UPLC® M-class, Peptide CSH™ 130A, 1.7µm 75µm x 250mm, Waters Corp, Milford, MA, USA) at 300 nL/min with a gradient from 5-22% over 75 min. The LC aqueous mobile phase was 0.1% (v/v) formic acid in water and the organic mobile phase was 0.1% (v/v) formic acid in 100% (v/v) acetonitrile. Mass spectra were acquired using the data-dependent mode with a mass range of m/z 375–1500, resolution 120,000, AGC (automatic gain control) target 4 x 10 5, maximum injection time 50 ms for the MS1. Data-dependent MS2 spectra were acquired by HCD in the ion trap with a normalized collision energy (NCE) set at 30%, AGC target set to 5 x 10 4 and a maximum injection time of 86 ms. The identification and quantitation of the proteins were done using Proteome Discoverer (Version 2.4; Thermo Fisher Scientific) utilizing MASCOT search engine (Version 2.7.0; Matrix Science Ltd, London, UK)^56^. The search was performed against an in-house modified version of the cRAP database (theGPM.org/cRAP) and the *Rhodopseudomonas palustris* (version_20230110) database obtained from UniProt (ID: UP000001426_258594, www.uniprot.org), assuming the digestion enzyme trypsin and a maximum of 2 missed cleavages. Mascot was searched with a fragment ion mass tolerance of 0.06 Da and a parent ion tolerance of 15.0 ppm. Deamidated of asparagine and glutamine, oxidation of methionine, were specified in Mascot as variable modifications while carbamidomethylation of cysteine was fixed. Peptides were validated by Percolator with a 0.01 posterior error probability (PEP) threshold. The data were searched using a decoy database to set the false discovery rate (FDR) to 1% (high confidence). Only proteins identified with a minimum of 2 unique peptides and 5 peptide-spectrum matches (PSMs) were further analyzed for quantitative changes. The peptides were quantified using the precursor abundance based on intensity. The peak abundance was normalized using total peptide amount. The peptide group abundances are summed for each sample and the maximum sum for all files is determined. The normalization factor used is the factor of the sum of the sample and the maximum sum in all files. The normalized abundances were scaled for each protein so that the average abundance is 100. The mass spectrometry proteomics data have been deposited at the ProteomeXchange Consortium via the PRIDE 37 partner repository with the dataset identifier PXDXXXXX.

### Metabolite Analysis

To obtain metabolite profiles, cell pellets were suspended into 1.0 mL of extraction solution (80% MeOH, 20%H2O, containing 2.13uM 13C5-15N1-proline and 13C6-glucose) on Eppendorf tubes, on dry-ice. About 100 mg of ZrBO5 beads were added to each sample. The cells were disrupted using a Bullet Blender-24 at 4 oC, using a setting of 7 for 3 min. The disruption cycle was repeated 2 times. The samples were then centrifuged at 4 oC, at 15,000xg for 10 min. The supernatants were decanted into clean Eppendorf tubes and the solvent was evaporated overnight, also at 4 oC. The pellets were suspended into 50 uL of LC-MS grade water and transferred into HPLC V-vials and kept at 5 oC in the autosampler. An Agilent LC-1200 HPLC system was used to stored and separate the compounds. The mobile phase consisted of 0.1% formic acid in water (A) and acetonitrile (B). A 2.1×100 mm Cortecs C18 Reverse Phase column (Millipore, Milford, MA) was used to separate the compounds. The flow rate was set at 0.25 mL/min. The effluent was continuously fed to a 4000 QTrap (Sciex, Framingham, MA) operating in multiple reaction monitoring mode (MRM). The gradient program was as follows: 0 to 2 min, 95%A, 2 to 7 min, 95% à 5% A, 14 to 15 min, 5% A, 15 to 20 min, 95% A. The instrument operated in negative ionization mode (Turbo V ion source), with the following parameters: IS= −4000V; CUR= 20 psi (N2); T = 600 oC, GS1= 70 psi; GS2=30 psi, DP= −50 V, EP=-10 V, CXP=-10 V.

### Omics data analysis and statistical methods

Differential Gene Expression analysis on transcriptome profiles obtained from CD genomics was performed using the DESeq2 R package in RStudio. Differentially abundant proteins were obtained from proteomic profiles using Perseus Software. Protein abundances that were not detected in 3 of 5 samples in each sample group were discarded. Abundances were then quantile normalized, and Log_2_ transformed before imputing missing values. *p-*values were then obtained and adjusted using Benjamini-Hochberg method and Log_2_ Fold Change values were obtained. t-SNE plots were obtained via Python.

## Supporting information

Supplementary File description

## Author Contributions

MK, NBC, RS, CI, and AA contributed in order of prevalence to this work. MK and CI produced wild type growth curves. AA performed a comparative genomics analysis for determining ligninolysis pathways. MK, CI, and DM harvested cultures for obtaining the transcriptomic dataset while MK harvested cultures for the proteomic and metabolite datasets. MK developed methodology, NBC provided metabolic modelling predictions, CI developed the CRISPRi expression vector and MK designed vectors for repressing select *R. palustris* transcripts. MK wrote the manuscript with contribution from NBC and guidance from RS and CI.

## Acknowledgements

We gratefully acknowledge funding support from National Science Foundation (NSF) CAREER grant (25-1106-0039-001) awarded to R.S. We thank the Proteomics & Metabolomics Facility (RRID:SCR_021314), Nebraska Center for Biotechnology at the University of Nebraska-Lincoln for the mass spectrometry analysis as well as Javier Seravalli from the UNL Spectroscopy and Biophysics Core for providing metabolite profiles. The facility and instrumentation are supported by the Nebraska Research Initiative. We would also like to thank CD Genomics for generating the corresponding transcriptomic dataset.

