## Supplementary File description for "High Enzyme Promiscuity in Lignin Degradation Mechanisms in *Rhodopseudomonas palustris* CGA009"

Description of additional supplementary files

Title: Supplementary Data 1

Description: Raw data for all depicted R. palustris growth curves.

Title: Supplementary Data 2.

Description: Comparative expression data generated by DEseq2.

Title: Supplementary Data 3.

Description: Comparative protein abundance data generated by Perseus.

Title: Supplementary Data 4.

Description: Metabolite profiles of aerobic and anaerobic R. palustris cultures grown on various LBPs.

Title: Supplementary Data 5.

Description: Redox-homeostasis modelling results.
