## Supplementary File for "High Enzyme Promiscuity in Lignin Degradation Mechanisms in *Rhodopseudomonas palustris* CGA009"

**Supplementary Table I.** Plasmids used in this work.

| **Name** | **Parts** | **Type and Source** |
| --- | --- | --- |
| BBR1-kan-LacI | pBBR1 replicon; *kan^R^; P_Lac_:lacI,* | Replicative  This Study |
| BBR1-kan-dCas9-ControlVector | pBBR1 replicon; *kan^R^; P_Lac_:lacI, dCas9, rrnB, T7Te, J23119, sgRNA scaffold, tonB* | Replicative  This Study |
| BBR1-kan-dCas9- RPA0650 | pBBR1 replicon; *kan^R^; P_Lac_:lacI, dCas9, rrnB, T7Te, J23119, RPA0650sgRNA*, *sgRNA scaffold, tonB* | Replicative  This Study |
| BBR1-kan-dCas9- RPA1206 | pBBR1 replicon; *kan^R^; P_Lac_:lacI, dCas9, rrnB, T7Te, J23119, RPA1206sgRNA, sgRNA scaffold, tonB* | Replicative  This Study |
| BBR1-kan-dCas9- RPA1580 | pBBR1 replicon; *kan^R^; P_Lac_:lacI, dCas9, rrnB, T7Te, J23119, RPA1580sgRNA, sgRNA scaffold, tonB* | Replicative  This Study |
| BBR1-kan-dCas9- RPA1786 | pBBR1 replicon; *kan^R^; P_Lac_:lacI, dCas9, rrnB, T7Te, J23119, RPA1786sgRNA, sgRNA scaffold, tonB* | Replicative  This Study |
| BBR1-kan-dCas9- RPA1787 | pBBR1 replicon; *kan^R^; P_Lac_:lacI, dCas9, rrnB, T7Te, J23119, RPA1787sgRNA, sgRNA scaffold, tonB* | Replicative  This Study |
| BBR1-kan-dCas9- RPA3701 | pBBR1 replicon; *kan^R^; P_Lac_:lacI, dCas9, rrnB, T7Te, J23119, RPA3701sgRNA, sgRNA scaffold, tonB* | Replicative  This Study |
| BBR1-kan-dCas9- RPA3717 | pBBR1 replicon; *kan^R^; P_Lac_:lacI, dCas9, rrnB, T7Te, J23119, RPA3717sgRNA, sgRNA scaffold, tonB* | Replicative  This Study |
| BBR1-kan-dCas9- RPA4308 | pBBR1 replicon; *kan^R^; P_Lac_:lacI, dCas9, rrnB, T7Te, J23119, RPA4308sgRNA, sgRNA scaffold, tonB* | Replicative  This Study |
| BBR1-kan-dCas9- RPA4696 | pBBR1 replicon; *kan^R^; P_Lac_:lacI, dCas9, rrnB, T7Te, J23119, RPA4696sgRNA, sgRNA scaffold, tonB* | Replicative  This Study |

**Supplementary Table II.**  List of genetic parts used in this work^a^.

| Part name | Type and source | DNA sequence |
| --- | --- | --- |
| *kan^R^* | Antibiotic resistance gene  (Kovach et al., 1995) | atgattgaacaagatggattgcacgcaggttctccggccgcttgggtggagaggctattcggctatgactgggcacaacagacaatcggctgctctgatgccgccgtgttccggctgtcagcgcaggggcgcccggttctttttgtcaagaccgacctgtccggtgccctgaatgaactgcaggacgaggcagcgcggctatcgtggctggccacgacgggcgttccttgcgcagctgtgctcgacgttgtcactgaagcgggaagggactggctgctattgggcgaagtgccggggcaggatctcctgtcatctcaccttgctcctgccgagaaagtatccatcatggctgatgcaatgcggcggctgcatacgcttgatccggctacctgcccattcgaccaccaagcgaaacatcgcatcgagcgagcacgtactcggatggaagccggtcttgtcgatcaggatgatctggacgaagagcatcaggggctcgcgccagccgaactgttcgccaggctcaaggcgcgcatgcccgacggcgaggatctcgtcgtgacccatggcgatgcctgcttgccgaatatcatggtggaaaatggccgcttttctggattcatcgactgtggccggctgggtgtggcggaccgctatcaggacatagcgttggctacccgtgatattgctgaagagcttggcggcgaatgggctgaccgcttcctcgtgctttacggtatcgccgctcccgattcgcagcgcatcgccttctatcgccttcttgacgagttcttctga |
| *P_Lac_* | Promoter  (Kovach et al., 1995) | ggcagtgagcgcaacgcaattaatgtgagttagctcactcattaggcaccccaggctttacactttatgcttccggctcgtatgttgtgtggaattgtgagcggataacaat |
| *dCas9* | AddGene | atggataagaaatactcaataggcttagctatcggcacaaatagcgtcggatgggcggtgatcactgatgaatataaggttccgtctaaaaagttcaaggttctggaaaatacagaccgccacagtatcaaaaaaaatcttataggggctcttttatttgacagtggagagacagcggaagcgactcgtctcaaacggacagctcgtagaaggtatacacgtcggaagaatcgtatttgttatctacaggagattttttcaaatgagatggcgaaagtagatgatagtttctttcatcgacttgaagagtcttttttggtggaagaagacaagaagcatgaacgtcatcctatttttggaaatatagtagatgaagttgcttatcatgagaaatatccaactatctatcatctgcgaaaaaaattggtagattctactgataaagcggatttgcgcttaatctatttggccttagcgcatatgattaagtttcgtggtcattttttgattgagggagatttaaatcctgataatagtgatgtggacaaactatttatccagttggtacaaacctacaatcaattatttgaagaaaaccctattaacgcaagtggagtagatgctaaagcgattctttctgcacgattgagtaaatcaagacgattagaaaatctcattgctcagctccccggtgagaagaaaaatggcttatttgggaatctcattgctttgtcattgggtttgacccctaattttaaatcaaattttgatttggcagaagatgctaaattacagctttcaaaagatacttacgatgatgatttagataatttattggcgcaaattggagatcaatatgctgatttgtttttggcagctaagaatttatcagatgctattttactttcagatatcctaagagtaaatactgaaataactaaggctcccctatcagcttcaatgattaaacgctacgatgaacatcatcaagacttgactcttttaaaagctttagttcgacaacaacttccagaaaagtataaagaaatcttttttgatcaatcaaaaaacggatatgcaggttatattgatgggggagctagccaagaagaattttataaatttatcaaaccaattttagaaaaaatggatggtactgaggaattattggtgaaactaaatcgtgaagatttgctgcgcaagcaacggacctttgacaacggctctattccccatcaaattcacttgggtgagctgcatgctattttgagaagacaagaagacttttatccatttttaaaagacaatcgtgagaagattgaaaaaatcttgacttttcgaattccttattatgttggtccattggcgcgtggcaatagtcgttttgcatggatgactcggaagtctgaagaaacaattaccccatggaattttgaagaagttgtcgataaaggtgcttcagctcaatcatttattgaacgcatgacaaactttgataaaaatcttccaaatgaaaaagtactaccaaaacatagtttgctttatgagtattttacggtttataacgaattgacaaaggtcaaatatgttactgaaggaatgcgaaaaccagcatttctttcaggtgaacagaagaaagccattgttgatttactcttcaaaacaaatcgaaaagtaaccgttaagcaattaaaagaagattatttcaaaaaaatagaatgttttgatagtgttgaaatttcaggagttgaagatagatttaatgcttcattaggtacctaccatgatttgctaaaaattattaaagataaagattttttggataatgaagaaaatgaagatatcttagaggatattgttttaacattgaccttatttgaagatagggagatgattgaggaaagacttaaaacatatgctcacctctttgatgataaggtgatgaaacagcttaaacgtcgccgttatactggttggggacgtttgtctcgaaaattgattaatggtattagggataagcaatctggcaaaacaatattagattttttgaaatcagatggttttgccaatcgcaattttatgcagctgatccatgatgatagtttgacatttaaagaagacattcaaaaagcacaagtgtctggacaaggcgatagtttacatgaacatattgcaaatttagctggtagccctgctattaaaaaaggtattttacagactgtaaaagttgttgatgaattggtcaaagtaatggggcggcataagccagaaaatatcgttattgaaatggcacgtgaaaatcagacaactcaaaagggccagaaaaattcgcgagagcgtatgaaacgaatcgaagaaggtatcaaagaattaggaagtcagattcttaaagagcatcctgttgaaaatactcaattgcaaaatgaaaagctctatctctattatctccaaaatggaagagacatgtatgtggaccaagaattagatattaatcgtttaagtgattatgatgtcgatgccattgttccacaaagtttccttaaagacgattcaatagacaataaggtcttaacgcgttctgataaaaatcgtggtaaatcggataacgttccaagtgaagaagtagtcaaaaagatgaaaaactattggagacaacttctaaacgccaagttaatcactcaacgtaagtttgataatttaacgaaagctgaacgtggaggtttgagtgaacttgataaagctggttttatcaaacgccaattggttgaaactcgccaaatcactaagcatgtggcacaaattttggatagtcgcatgaatactaaatacgatgaaaatgataaacttattcgagaggttaaagtgattaccttaaaatctaaattagtttctgacttccgaaaagatttccaattctataaagtacgtgagattaacaattaccatcatgcccatgatgcgtatctaaatgccgtcgttggaactgctttgattaagaaatatccaaaacttgaatcggagtttgtctatggtgattataaagtttatgatgttcgtaaaatgattgctaagtctgagcaagaaataggcaaagcaaccgcaaaatatttcttttactctaatatcatgaacttcttcaaaacagaaattacacttgcaaatggagagattcgcaaacgccctctaatcgaaactaatggggaaactggagaaattgtctgggataaagggcgagattttgccacagtgcgcaaagtattgtccatgccccaagtcaatattgtcaagaaaacagaagtacagacaggcggattctccaaggagtcaattttaccaaaaagaaattcggacaagcttattgctcgtaaaaaagactgggatccaaaaaaatatggtggttttgatagtccaacggtagcttattcagtcctagtggttgctaaggtggaaaaagggaaatcgaagaagttaaaatccgttaaagagttactagggatcacaattatggaaagaagttcctttgaaaaaaatccgattgactttttagaagctaaaggatataaggaagttaaaaaagacttaatcattaaactacctaaatatagtctttttgagttagaaaacggtcgtaaacggatgctggctagtgccggagaattacaaaaaggaaatgagctggctctgccaagcaaatatgtgaattttttatatttagctagtcattatgaaaagttgaagggtagtccagaagataacgaacaaaaacaattgtttgtggagcagcataagcattatttagatgagattattgagcaaatcagtgaattttctaagcgtgttattttagcagatgccaatttagataaagttcttagtgcatataacaaacatagagacaaaccaatacgtgaacaagcagaaaatattattcatttatttacgttgacgaatcttggagctcccgctgcttttaaatattttgatacaacaattgatcgtaaacgatatacgtctacaaaagaagttttagatgccactcttatccatcaatccatcactggtctttatgaaacacgcattgatttgagtcagctaggaggtgactga |
| *pLacI* | AddGene | attcaccaccctgaattgactctcttccgggcgctatcatgccataccgcgaaaggttttgcgccattcgatggtgtc |
| *LacI* | AddGene | tcactgcccgctttccagtcgggaaacctgtcgtgccagctgcattaatgaatcggccaatgcgcggggagaggcggtttgcgtattgggcgccagggtggtttttcttttcaccagtgagacgggcaacagctgattgcccttcaccgcctggccctgagagagttgcagcaagcggtccacgctggtttgccccagcaggcgaaaatcctgtttgatggtggttaacggcgggatataacatgagctgtcttcggtatcgtcgtatcccactaccgagatgtccgcaccaacgcgcagcccggactcggtaatggcgcgcattgcgcccagcgccatctgatcgttggcaaccagcatcgcagtgggaacgatgccctcattcagcatttgcatggtttgttgaaaaccggacatggcactccagtcgccttcccgttccgctatcggctgaatttgattgcgagtgagatatttatgccagccagccagacgcagacgcgccgagacagaacttaatgggcccgctaacagcgcgatttgctggtgacccaatgcgaccagatgctccacgcccagtcgcgtaccgtcttcatgggagaaaataatactgttgatgggtgtctggtcagagacatcaagaaataacgccggaacattagtgcaggcagcttccacagcaatggcatcctggtcatccagcggatagttaatgatcagcccactgacgcgttgcgcgagaagattgtgcaccgccgctttacaggcttcgacgccgcttcgttctaccatcgacaccaccacgctggcacccagttgatcggcgcgagatttaatcgccgcgacaatttgcgacggcgcgtgcagggccagactggaggtggcaacgccaatcagcaacgactgtttgcccgccagttgttgtgccacgcggttgggaatgtaattcagctccgccatcgccgcttccactttttcccgcgttttcgcagaaacgtggctggcctggttcaccacgcgggaaacggtctgataagagacaccggcatactctgcgacatcgtataacgttactggtttcac |
| *pLac* | AddGene | tttacactttatgcttccggctcgtatgttg |
| *Lac operator* | AddGene | ttgtgagcggataacaa |
| *rrnB* | AddGene | caaataaaacgaaaggctcagtcgaaagactgggcctttcgttttatctgttgtttgtcggtgaacgctctc |
| *T7Te* | AddGene | ggctcaccttcgggtgggcctttctgcg |
| *J23119* | AddGene | ttgacagctagctcagtcctaggtataatgctagc |
| *sgRNA scaffold* | AddGene | gttttagagctagaaatagcaagttaaaataaggctagtccgttatcaacttgaaaaagtggcaccgagtcggtgc |
| *tonB* | AddGene | agtcaaaagcctccgaccggaggcttttgact |

^a^ <http://www.ncbi.nlm.nih.gov/gene>

**Supplementary Table III.**  sgRNA sequences for targeted dCas9 suppression^a^.

| Primer Name | DNA Sequence | Predicted Suppression | | | Log_2_ Fold Repression | Purpose |
| --- | --- | --- | --- | --- | --- | --- |
| *RPA0650* | CCTATCCTCACCGAAACCCAGGG | | 67.01% | 5.78 | | Redox |
| *RPA1206* | CCAGATGCGGATCTTCCAGGAGG | | 73.93% | nd | | Catabolism |
| *RPA1580* | CCGTGCCTCAGAGCTCACCGAGG | | 78.66% | 3.815 | | Redox |
| *RPA1786* | GCGCCCTGTCAGCATCATGTCGG | | 69.57% | 2.88 | | Catabolism |
| *RPA3701* | CTTTTACACCATGAACCGCGCGG | | 72.61% | -0.26 | | Redox |
| *RPA3717* | GTGTACCCGATGATCAACGAGGG | | 75.26% | 2.14 | | Redox |
| *RPA4308* | ATCCTTGAAGATCTGCACGGCGG | | 79.52% | 1.99 | | Redox |
| *RPA4696* | TCGGTGCGAAACATCACCAGCGG | | 75.60% | 1.06 | | Catabolism |

^a^ https://chopchop.cbu.uib.no/

**Supplementary Table VI.** RT-qPCR primers.

| **Primer**  **Name** | **Sequence** | **Amplicon Length (bp)** | **Primer Concentration (nM)** | **cDNA or gDNA Concentration (ng/μL)** | **Primer**  **Efficiency**  **(%)** | **R^2** |
| --- | --- | --- | --- | --- | --- | --- |
| 16SrRNA F | GCTTAACACAT  GCAAGTCGAAC | 230 | 40 | 0.05 | 98.61 | 0.994 |
| 16SrRNA R2 | TCATCCTCTCA  GACCAGCTAC |  |  |  |  |  |
| *RPA0650 F3* | GATCGCTATGGTCTGGTCTCC | 128 | 50 | 100 | 102.26 | 0.989 |
| *RPA0650 R* | CGATTGAGGCTTCCTTCAGC |  |  |  |  |  |
| *RPA1206 F* | GCTCGACAAGATCCTGTCCTACATC | 169 | 50 | 100 | nd | NA |
| *RPA1206 R* | GAAGATCTCCTCCTGGAAGATCC |  |  |  |  |  |
| *RPA1580 F3* | CTGATCGGCAATCACTTCGTG | 101 | 50 | 100 | 99.15 | 0.977 |
| *RPA1580 R3* | AGCATGTCGTAAGAATAGCGAGAGC |  |  |  |  |  |
| *RPA1786 F2* | CTACAGGCGTTGCCGATGATC | 136 | 50 | 100 | 106.28 | 0.983 |
| *RPA1786 R* | TGGCTGTCTTGTGATCGAGGAAAG |  |  |  |  |  |
| *RPA3701 F3* | CATCACCCAGTTCTTCTTCGATAACG | 119 | 50 | 100 | 100.15 | 0.997 |
| *RPA3701 R* | CCTGCTTGAAGTTCTGCACC |  |  |  |  |  |
| *RPA3717 F* | GTCGAACACCTCGTATCTGTCGATC | 115 | 50 | 100 | 101.35 | 0.986 |
| *RPA3717 R2* | GATTTCGCACAGCTTCATCACGTTG |  |  |  |  |  |
| *RPA4308 F* | GTTCGACGTGTTCTCTGAAGAGC | 153 | 50 | 100 | 102.21 | 0.984 |
| *RPA4308 R* | GTCAGCAGATAGTCGCTCATCTG |  |  |  |  |  |
| *RPA4696 F2* | GAGGTGACCTGCATCGACAAC | 136 | 50 | 100 | 97.712 | 1 |
| *RPA4696 R* | CGATCCGCAGTTTTTCCAGC |  |  |  |  |  |

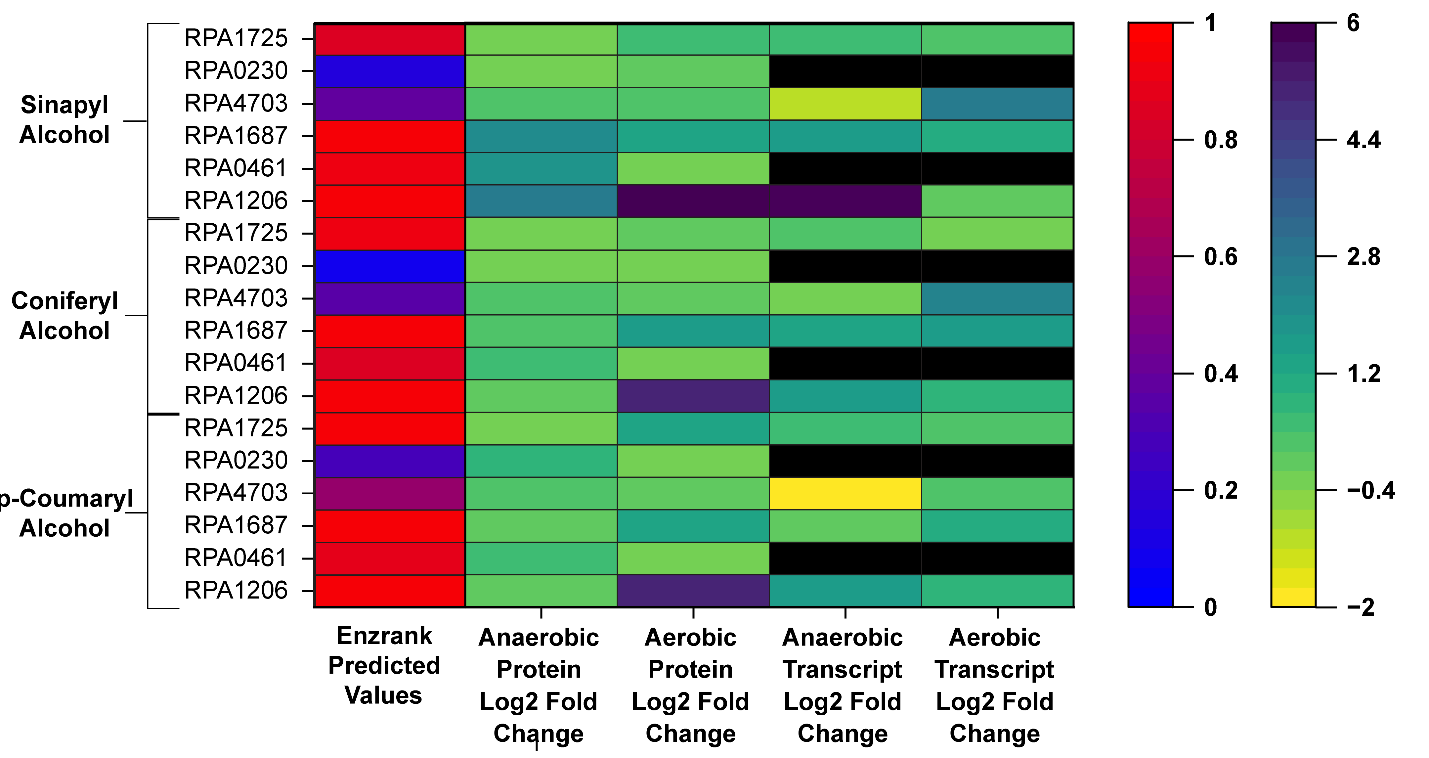

**Supplementary Figure 1.** Combined heatmap for aldehyde dehydrogenase EnzRank predictions for multiple upregulated aldehyde dehydrogenases. Predictions and upregulation states are depicted for *R. palustris* cultures grown under each type of lignin both aerobically and anaerobically.

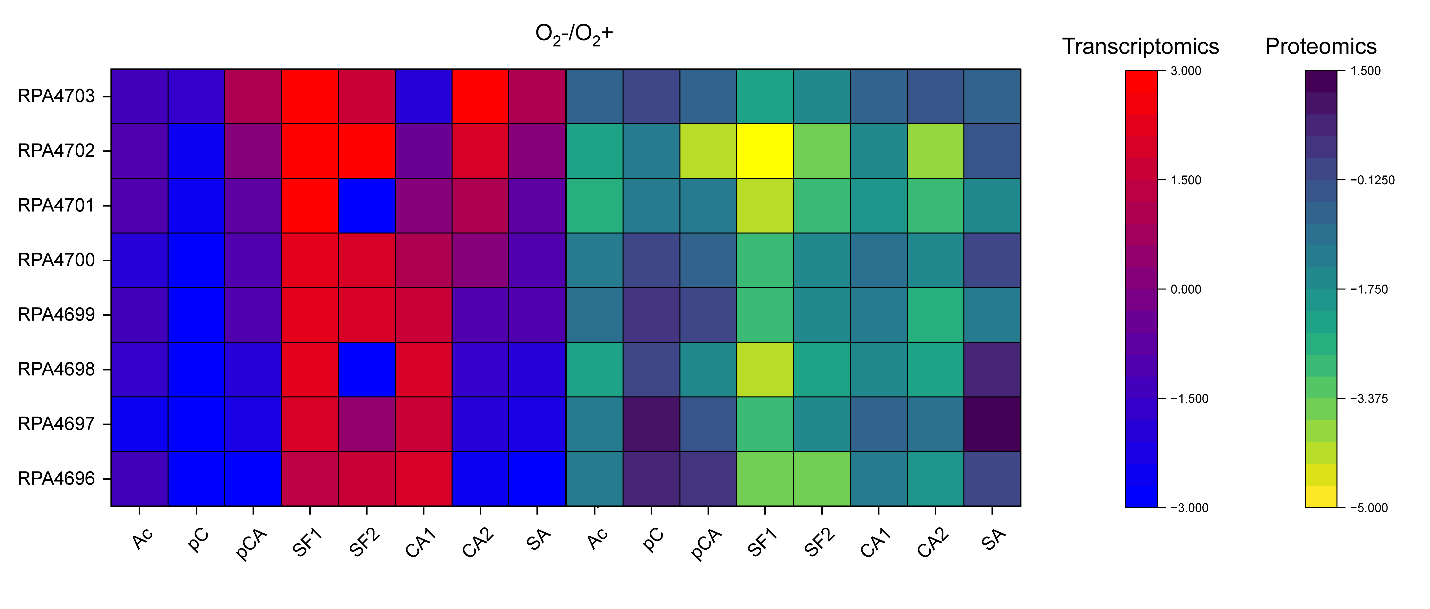

**Supplementary Figure 2.** Upregulation heatmap for meta-cleavage of anaerobic state compared to corresponding aerobic state. SF1 represents sodium ferulate only samples, while SF2 represents sodium ferulate and acetate supplemented samples. CA1 represents coniferyl alcohol only samples, while CA2 represents coniferyl alcohol and acetate supplemented samples. Meta-cleavage is seemingly upregulated under anaerobic conditions in some samples.

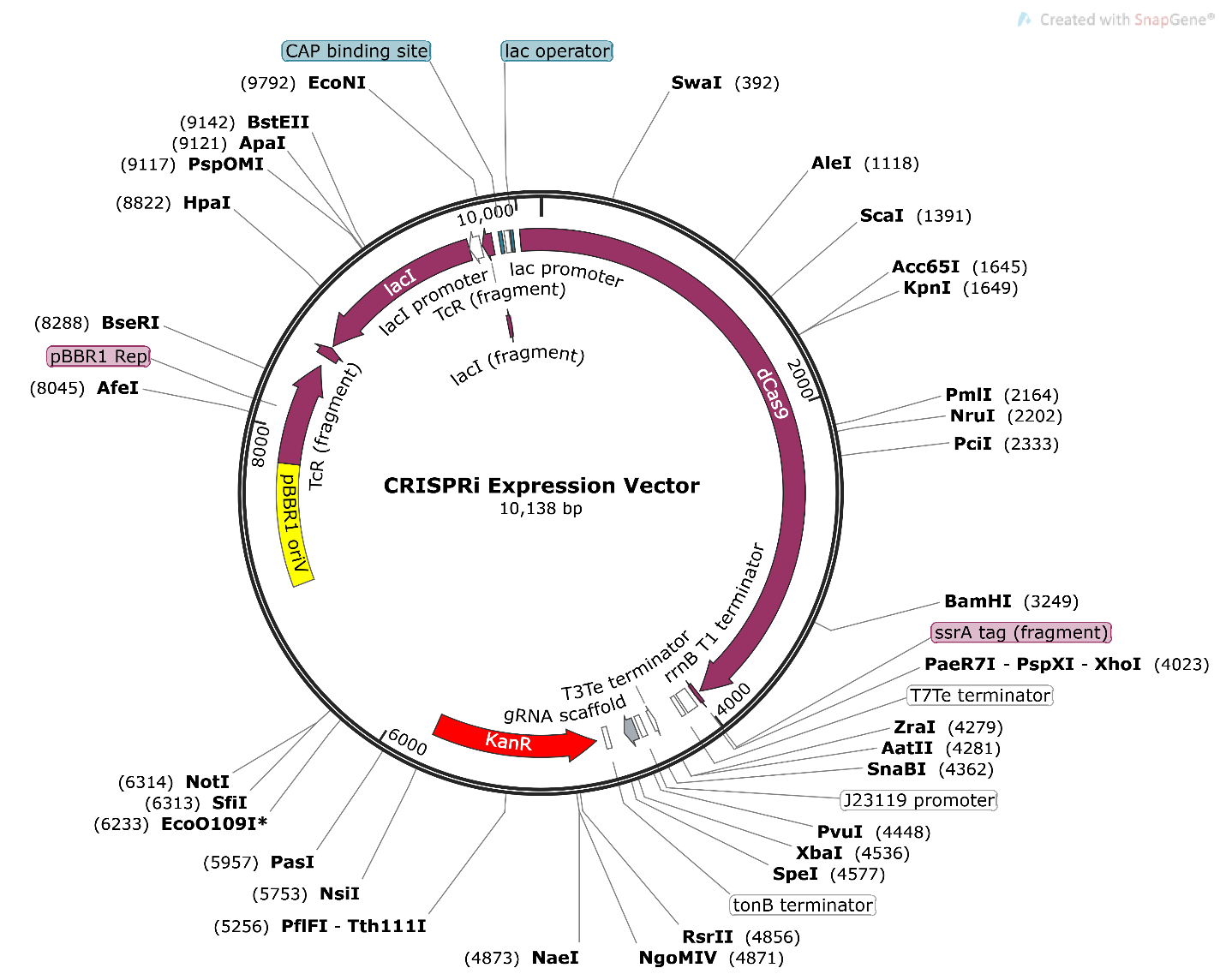

**Supplementary Figure 3.** Map of dCas9 expression vector with interchangeable sgRNA sequences.
